# M102, a combined NRF2 and HSF-1 activator for neuroprotection in amyotrophic lateral sclerosis

**DOI:** 10.1101/2024.12.08.627389

**Authors:** Amy F. Keerie, Raquel Rua Martins, Chloe F. Allen, Katie Bowden, Sufana Al Mashhadi, Thomas Marlow, Monika Myszczynska, Nikitha Thakur, Selina N. Beal, Allan Shaw, Shivani Suresh, Scott N. McKinnon, Alex Daniel, Tyler Wells, Ira N. Kalfus, Ning Shan, Pamela J. Shaw, Laura Ferraiuolo, Richard J. Mead

## Abstract

M102 is a central nervous system (CNS) penetrant small molecule electrophile which activates *in vivo* the NFE2-related factor 2 antioxidant response element (NRF2-ARE) pathway, as well as transcription of heat-shock element (HSE) associated genes. In the TDP-43^Q331K^ transgenic mouse model of ALS dosed subcutaneously at 5mg/kg OD or 2.5mg/kg BD with M102, significant improvements in compound muscle action potential (CMAP) amplitude of hind limb muscles and gait parameters were observed at 6 months of age, with associated target engagement.

An oral dose response study of M102 in SOD1^G93A^ transgenic mice showed a dose-dependent improvement in the CMAP of hindlimb muscles which correlated with preservation of lumbar spinal motor neurons at the same time point. These data enabled prediction of human efficacious exposures and doses, which were well within the safety margin predicted from Good Laboratory Practice (GLP) toxicology studies.

A parallel program of work *in vitro* showed that M102 rescued motor neuron survival in co-culture with patient-derived astrocytes from sporadic, *C9orf72* and *SOD1* cases. Markers of oxidative stress, as well as indices of TDP-43 proteinopathy were also reduced by exposure to M102 in these *in vitro* models.

This comprehensive package of preclinical efficacy data across two mouse models as well as patient-derived astrocyte toxicity assays, provides a strong rationale for clinical evaluation of M102 in ALS patients. Combined with the development of target engagement biomarkers and the completed preclinical toxicology package, a clear translational pathway to testing in ALS patients has been developed.

**One Sentence Summary:** M102, a dual NRF2/HSF1 activator, slows disease progression in 2 mouse models and exerts neuroprotection in human cellular models of ALS.

## INTRODUCTION

Amyotrophic lateral sclerosis (ALS) is a rapidly progressive, fatal neurodegenerative disorder in which motor neuron injury and cell death causes muscle weakness and wasting, leading to progressive loss of motor control of the upper and lower limbs as well as bulbar and respiratory functions. The lifetime risk is approximately 1 in 300-350 (*1*) and the average course of the disease is 2.5-3 years from symptom onset (*2*). Currently, there are no effective treatments to halt or reverse the progression of ALS and approved drugs only marginally increase survival (riluzole) or disease progression (edaravone) (*3–5*). The anti-sense oligonucleotide (ASO) treatment tofersen (QALSODY) has beneficial clinical effects and biomarker readouts, but applies only to the ∼2 percent of ALS patients who harbor a *SOD1* mutation (*6*). A key feature of ALS is the speed of progression. This poses huge problems of adjustment for affected individuals, an escalating burden on carers and families, and a challenge to those purchasers and providers of healthcare who are involved in meeting the variable, rapidly changing and complex care needs (*7, 8*).

The pathophysiology of ALS is complex. More than 30 genes are known to cause or contribute to motor neuron degeneration in ALS (*9*). Even in the presence of a mutation in a gene such as SOD1, the structure and function of which are well understood, it is recognised that a cascade of multiple pathophysiological mechanisms within both motor neurons and neighboring glial cells contribute to neurodegeneration (*9–11*). Data from disease model systems and human biosamples provide strong evidence for a role of redox imbalance, inflammation, mitochondrial dysfunction and altered proteostasis, as key drivers of the pathobiology of ALS (*12–16*). Therapies targeting individual pathways have failed in the clinic or shown marginal efficacy. In addition, few studies in ALS have shown target engagement for the proposed therapeutic agent in the clinic (*17*) with the exception of tofersen (*6*). There is clearly a huge unmet need for effective neuroprotective therapies to slow disease progression in ALS.

The nuclear factor erythroid 1-like 2 (NRF2) is a master regulator of the antioxidant response and activates the expression of over 1000 genes with cytoprotective properties (*18–20*). NRF2 protein levels are highly regulated through several different mechanisms including negative regulation at the protein level by Kelch-like ECH-associated protein 1 (KEAP1) (*21*). There is a body of evidence from both cellular and animal models and human biosamples that this cytoprotective NRF2 system is dysregulated in ALS (*22–27*).

We previously identified M102 (chemical name: S-(+)-10,11-dihydroxyaporphine) in a compound screen to discover blood brain barrier penetrant NRF2-ARE pathway activators (*28*). We reported that M102 enhanced glutathione (GSH) secretion from astrocytes in co-culture; protected neuromuscular junctions from denervation in *SOD1*^G93A^ mice; and slowed the decline in motor function when dosed at 5 mg/kg subcutaneously daily. M102 also reduced the elevated basal oxidative stress seen in fibroblasts from ALS cases (*28*). M102 (S[+]-apomorphine) is an enantiomer of R-apomorphine which is used as a dopamine agonist in Parkinson’s disease. M102 itself is a very weak dopamine antagonist (*29*) and its structure is consistent with known NRF2 activators predicted to function through modification of cysteine residues on KEAP1, thereby reducing the degradation of NRF2, increasing NRF2 translocation to the nucleus and upregulating the expression of multiple cytoprotective genes (*18, 20*). M102, which we now demonstrate activates both the NRF2 and HSF-1 transcription factor pathways, has the potential to modulate all four of the key drivers of neurodegeneration in ALS as highlighted above. By targeting multiple pathways, we expect to significantly increase the probability of neuroprotection of motor neurons and clinical success.

Lack of translation between mouse models and clinical trials has hampered ALS treatment discovery over the last 20 years, with many compounds showing potential in mouse models, but failing to show efficacy in clinical trials (*17, 30, 31*). Here, we have generated an encouraging pre-clinical data set demonstrating significant positive effects in the same behavioral outputs in two robust and reproducible ALS mouse models, as well as a preclinical toxicology package that predicts safe efficacious doses in humans. In addition, we provide evidence that M102 increases motor neuron survival in an *in vitro* model of patient-derived astrocyte toxicity (*10*) across multiple subtypes of ALS. M102 reduces toxicity from astrocytes derived from *C9orf72, SOD1* and sporadic ALS patients and results in neuroprotection of co-cultured motor neurons. Overall, to our knowledge, this is one of the most comprehensive packages of pre-clinical efficacy for therapy development in ALS.

## RESULTS

### M102 activates NRF2 and HSF-1 transcription factor pathways in vivo

We previously identified M102 in a screen for CNS penetrant NRF2 activating molecules. We demonstrated that M102 activated NRF2-directed transcription *in vitro* and in the CNS and was able to rescue motor deficits in the SOD1^G93A^ mouse model of ALS at a daily subcutaneous dose of 5 mg/kg (*28*). *In vitro* evidence showed that M102 was able to increase production of glutathione from astrocytes, thereby mediating neuroprotection to co-cultured motor neurons. In order to expand the preclinical validation to other genetic subtypes of ALS we set out to establish a dose response for NRF2 activation in the CNS as a precursor to investigating efficacy in a second, *TARDBP* mutant ALS mouse model for which we have validated readouts (*32*).

Electrophilic compounds such as M102 also have the potential to activate HSF-1 (heat shock factor 1) signaling pathways (*33, 34*) and since HSF-1 activation would address additional pathogenic mechanisms in ALS (*35*), we also sought to establish whether M102 was able to transcriptionally activate canonical HSF-1 targets *in vivo* as well as NRF2-related targets.

We dosed wild-type (WT) mice with M102 at 0.5, 1.5, 5 and 10 mg/kg SC, once daily for 7 days, and measured the transcriptional response in cerebral cortex tissue using RT-qPCR. **Fig. 1** shows the transcriptional response of NRF2 (**Fig. 1A**) and HSF-1 (**Fig. 1B**) regulated genes. Gene targets of both transcription factors showed a clear dose response, with 5mg/kg being optimal in most cases. This represents the first indication that M102 is also able to activate HSF-1 directed transcription i*n vivo,* with HSF-1 regulated genes including Hspa1a (HSP70), Hspa8, and Syn1 showing elevated transcriptional responses.

**Figure 1:**
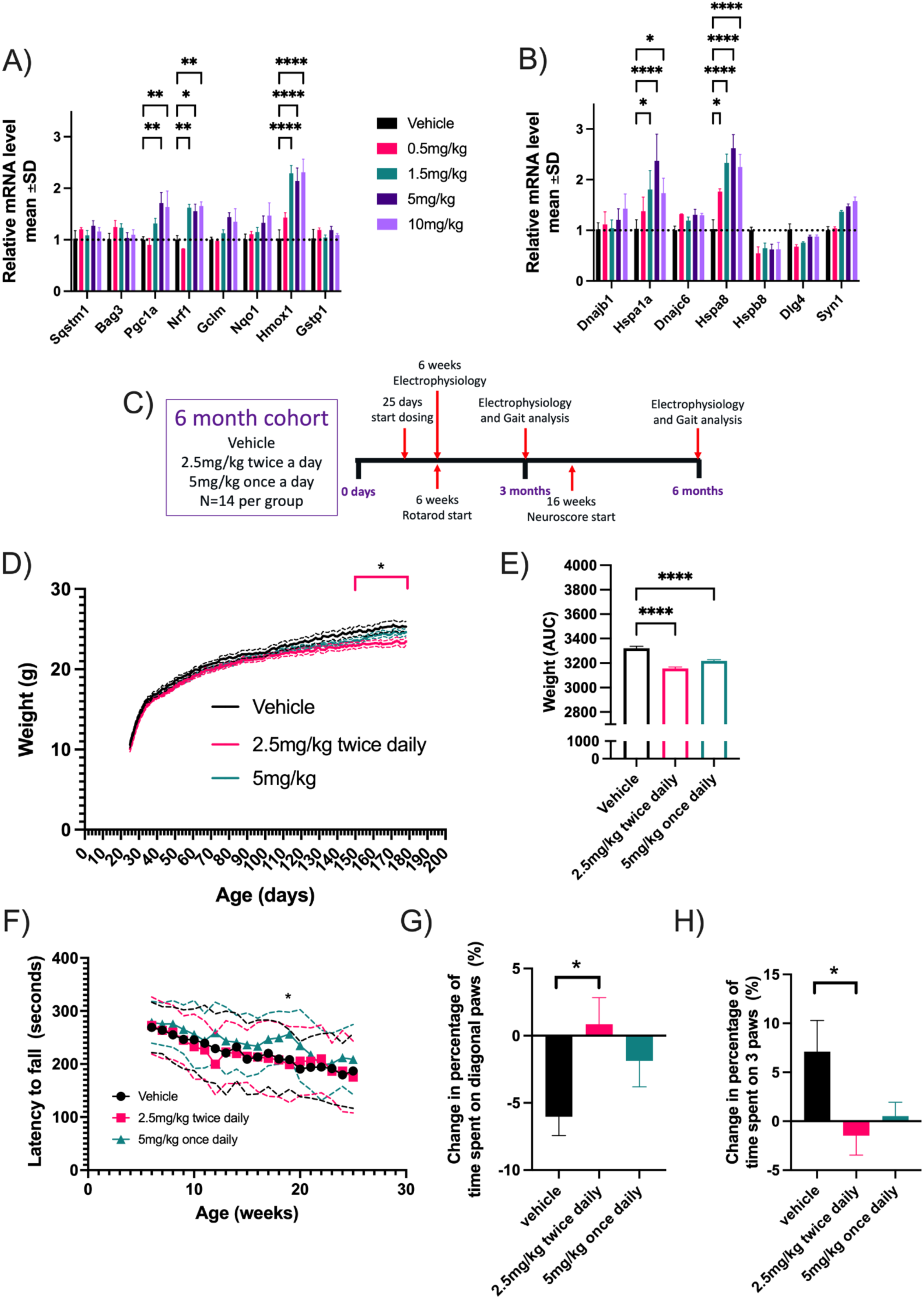
M102 treatment increases the activity of both NRF2 and HSF1 targets in WT mice and attenuates disease progression in TDP-43^Q331K^ mice. Changes in A) NRF2 and B) HSF1 targets 24 hours post dose after 7 days of dosing subcutaneously with different concentrations of M102 (N=3 per group). C) Diagram showing the design of the TDP-43^Q331K^ study with the major timepoints for electrophysiology and gait analysis. D) Body weight over time showing a reduction in the 2.5mg/kg BD M102 group when compared to vehicle dosed (N=13-14 per group). E) Area under the curve analysis of body weight showing reduction in both M102 dosed groups when compared to the vehicle dose group (N=13-14). F) Rotarod data shown as latency to fall (N=13-14 per group). Change in time spent on G) Diagonal paws and H) 3 paws showing significant improvement of the 2.5mg/kg BD M102 group when compared to the vehicle group (N=8 per group). All data shown as mean +/- SD. (* = p<0.05, ** = p<0.01, *** = p<0.001, **** = p<0.0001).

### M102 rescues motor, weight and, neurophysiological phenotypes in the TDP-43^Q331K^ transgenic mouse model of ALS

The study design for evaluating the effect of M102 in huTDP-43^Q331K^ transgenic mice is shown in **Fig. 1C**. Female TDP-43^Q331K^ mice were block randomised into three dosing groups and dosed subcutaneously with 5m/kg M102 as a split dose twice daily at 2.5mg/kg twice daily (BD) or a single 5mg/kg once daily (OD) dose from 25 days until 6 months of age. A variety of behavioral tests were performed at time points previously shown to correlate with significant milestones of disease progression, and CNS tissue was collected at 3 and 6 months.

Transgenic TDP-43^Q331K^ mice gain significantly more weight compared to their non-transgenic littermates (*32*). This is due to the increased amount of food they consume and the reduction in activity seen with this mutation in TDP-43 that is linked to an apathy fronto-temporal dementia (FTD) phenotype (*36*). Mouse weights were recorded daily before dosing to determine dose volume. **Fig. 1D** shows that weight gain is significantly reduced from 161 days in the 2.5mg/kg BD group (two way ANOVA with Dunnett’s post-test). At the end of the study there was a significant decrease in weight of the M102 2.5mg/kg BD dosed mice when compared to vehicle controls. At 177 days of age, M102 2.5mg/kg BD dosed animals weighed 23.2 +/- 1.9g compared to vehicle animals that weighed 25.4 +/- 2.3g (p = 0.05). When analysed as area under the curve, weight is significantly reduced in both 5mg/kg and 2.5 mg/kg BD dosed groups (**Fig. 1E**; one way ANOVA with Dunnett’s multiple cpmparison’s test p<0.05). Rotarod performance was also improved in the 5mg/kg dose group at 19 weeks of age (**Fig. 1F** - two way ANOVA with Dunnett’s post-testing).

Gait analysis was carried out at 3 and 6 months of age. An improvement in gait (**Fig. 1 G,H**) was observed in the 2.5mg/kg BD dosed group when compared to vehicle controls, using Catwalk (NoldusXT) gait analysis. This was shown as an increase in the amount of time spent using diagonal paws, a marker of normal mouse gait, versus a decrease in the amount of time spent on three paws, suggesting a reduction of gait unsteadiness (p<0.05 one way ANOVA, with Dunnett’s multiple comparison’s test).

Muscle electrophysiology was performed at 6 weeks and 6 months of age. At 6 weeks of age there was no significant difference in compound muscle action potential (CMAP) amplitude between either of the M102 dose groups and the vehicle dosed group. However, at 6 months of age there was a significant improvement in CMAP amplitude in both M102 treated groups when compared to the vehicle group (**Fig. 2 A**, p<0.01 at 2.5mg/kg BD and p<0.05 at 5mg/kg wo-way ANOVA with Dunnett’s multiple comparison’s test). The percentage change in CMAP amplitude between 6 weeks and 6 months of age also showed significant improvement of the M102 2.5mg/kg BD dosed group when compared to the vehicle dosed group (**Fig. 2B**, p<0.01 one-way ANOVA with Dunnet’s post test). At 6 months of age there was also a significant improvement in response to repetitive stimulation in the 2.5mg/kg BD dosed animals when compared to vehicle dosed animals, suggesting improved maintenance of the neuromuscular junctions (NMJs) (**Fig. 2C**, p<0.05 one-way ANOVA with Dunnett’s post test).

**Figure 2:**
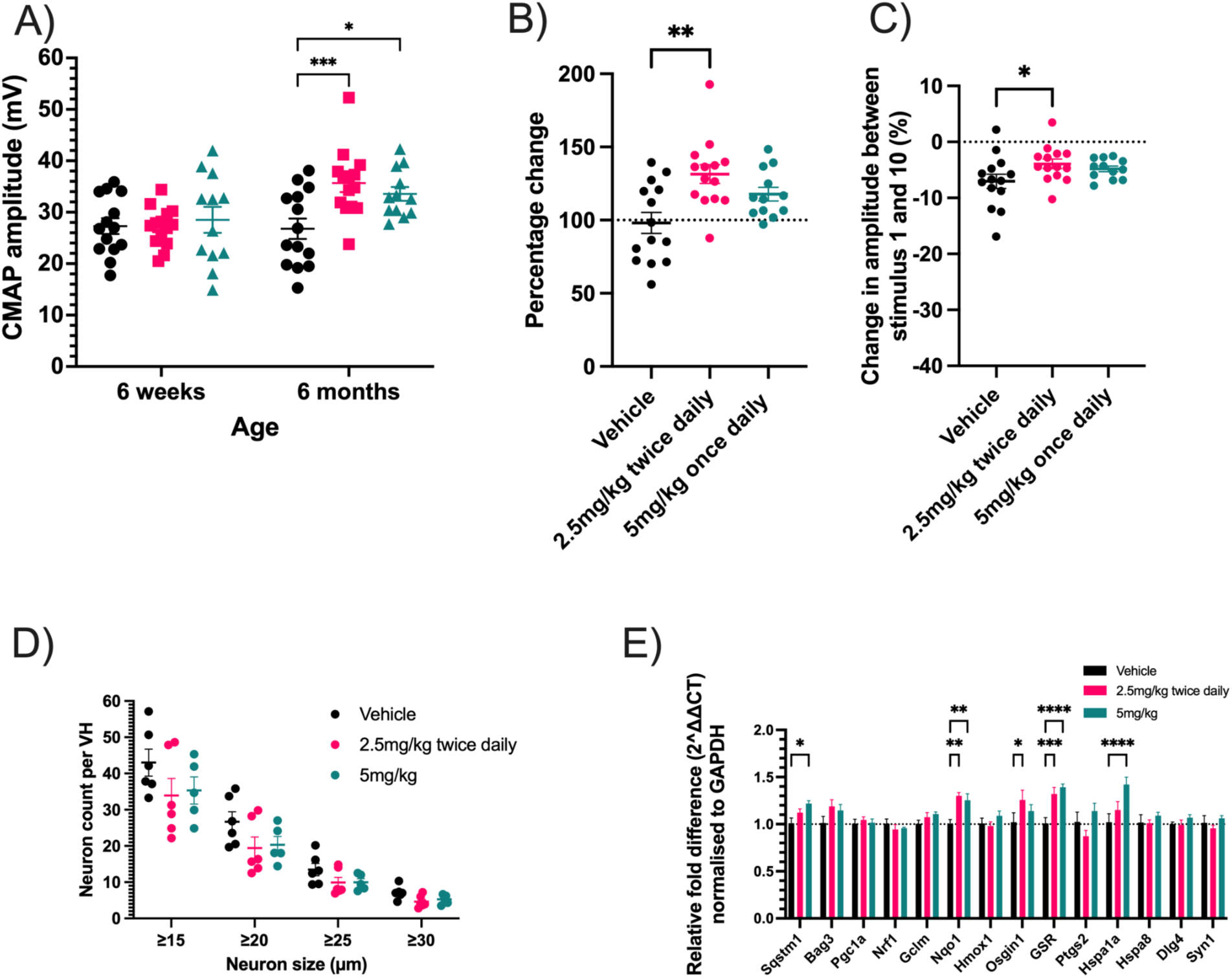
M102 treatment in TDP-43^Q331K^ mice improved electrophysiological performance by increasing NRF2 and HSF1 targets in the CNS. **A)** Compound muscle action potential (CMAP) amplitude of the hindlimb muscle at 6 weeks and 6 months of age and B) Percentage change in CMAP amplitude between 6 weeks and 6 months of age in M102 dosed groups compared to vehicle (N=12-14 per group). C) Change in repetitive stimulation amplitude between the 1st and 10th stimulus (N=12-14 per group). D) Motor neuron counts from lumbar spinal cord in size bins (N=5-6 per group). E) Cerebral cortex NRF2 and HSF1 targets after 3 months of dosing (N=6 per group). All data shown as mean +/- SD. (* = p<0.05, ** = p<0.01, *** = p<0.001, **** = p<0.0001).

Staining of lumbar spinal cords for Iba1 and GFAP showed no significant difference in the area stained or intensity of staining, suggesting that there was no change in microglial or astrocytic numbers in M102 dosed animals when compared to the vehicle group **(Suppl. Fig. 1A-F)**. Motor neuron counts and size distribution from the ventral horns of the lumbar spinal cord showed no significant difference between M102 dosed groups and vehicle dosed groups (**Fig. 2D**). However, in this model, we were unable to detect any significant difference between WT and TDP-43^Q331K^ transgenic mice in lumbar spinal cord motor neuron counts at a similar timepoint (**Suppl. Fig 2**).

Analysis of RNA levels in the cerebral cortex by RT-qPCR showed an increase in downstream targets of NRF2 and HSF-1 targets at 3 months of age in M102 dosed mice when compared to vehicle dosed mice, demonstrating activation of these pathways over time in the target tissue of the CNS (**Fig. 2E** two-way ANOVA followed by Dunnett’s multiple comparisons test).

### Pharmacokinetic data in wild-type mice (Suppl. Fig. 3)

Following a single subcutaneous dose of 5 mg/kg in mice, M102 exhibited a mean C_max_ (maximum plasma concentration) and AUC_last_ (area under the plasma concentration-time curve to the last measurable plasma concentration) of 989 ng/mL and 380 ng/mL*h, respectively. Pharmacokinetic (PK) evaluation of M102 post single dose oral administration was also performed. After a single oral dose of 10 mg/kg in mice, M102 showed a mean C_max_ and AUC_last_ of 168 ng/mL and 187 ng/mL*h, respectively. In the preclinical pharmacology studies in transgenic mouse models of ALS, M102 has demonstrated significant efficacy at a daily subcutaneous dose of 5 mg/kg (*28*). Assuming a correlation of M102 efficacy to its systemic exposure and dose proportionality, the efficacious oral dose of M102 was estimated to be 25 mg/kg or lower. This estimate was further confirmed in an additional *in vivo* pharmacology study in SOD1^G93A^ mice, where efficacy was observed at 12.5 and 25 mg/kg of M102 administered orally.

### Oral M102 SOD1^G93A^ study improves body weight, neurophysiology, and spinal cord motor neuron counts

We have previously shown disease modifying effects of M102 in the SOD1^G93A^ mouse model when M102 was dosed subcutaneously at 5mg/kg (*28*). In this study there was an upregulation of *Hmox1* and *Nqo1* in the spinal cord, a significant preservation of innervation at the neuromuscular junctions, a significant decrease in oxidised glutathione with an increase in the GSH/GSSG ratio in CNS tissue, and an improvement in rotarod performance and gait analysis parameters in the M102 exposed mice compared to the control group. We wanted to further expand on these data, by exploring the effect of M102 dosed orally at an equivalent dose to the subcutaneous dose, as this is the preferred route of administration in humans.

Transgenic female SOD1^G93A^ animals were block randomised into different groups and dosed orally with either vehicle, 5mg/kg, 12.5mg/kg or 25mg/kg M102 from 25 days until 90 days of age (n=8 per group). The doses were chosen as 5mg/kg subcutaneous dosing has an equivalent exposure compared to 25mg/kg oral dosing (**Suppl. Fig. 3**). Behavioral tests were carried out at various time points to determine motor function and tissue was collected at 90 days for histological and mRNA analysis (**Fig. 3A**).

**Figure 3:**
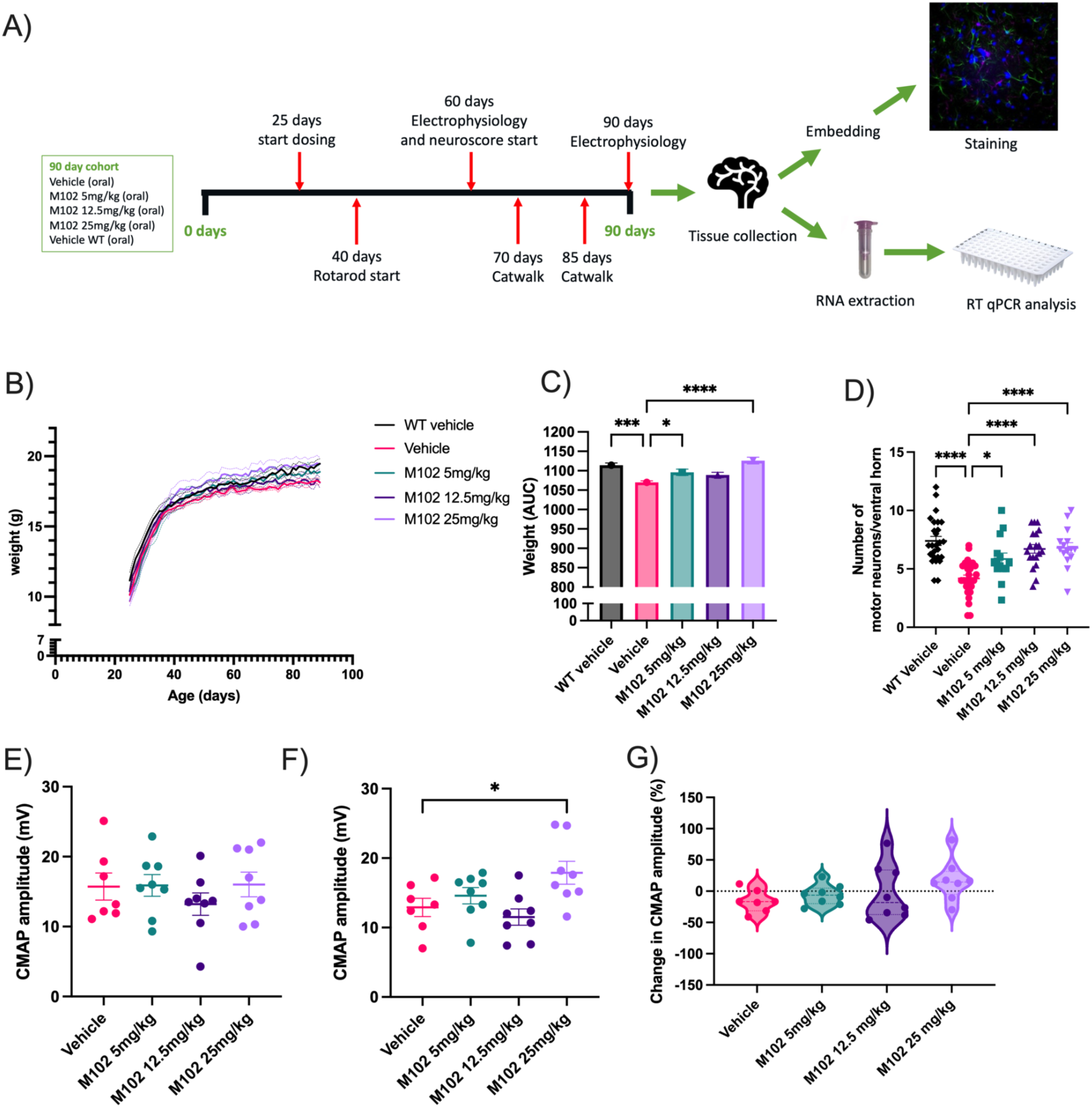
Oral M102 treatment in SOD1^G93A^ mice shows improvement in multiple parameters of disease progression. A) Design for the SOD1^G93A^ oral M102 study showing the main timepoints for electrophysiological and gait analysis. B) Body weights over time (N=8 per group). C) Area under the curve analysis of body weights (N=8 per group). D) Motor neuron counts per ventral horn (N=37-72 ventral horns per group). Compound muscle action potential (CMAP) amplitudes at E) 60 and F) 90 days of age (N= 7-8 per group). G) Percentage change in CMAP amplitude between 60 and 90 days of age (N=7-8 per group). All data shown as mean +/- SEM (* = p<0.05, ** = p<0.01, *** = p<0.001, **** = p<0.0001).

Animals were weighed daily before dosing to determine dose volume. SOD1^G93A^ transgenic mice are hypermetabolic and lose weight in the later stages of disease when compared to non-transgenic mice due to muscle and fat loss (*37, 38*). Area under the curve analysis showed a significant increase in body weight, and thus improvement of disease progression, of the 5mg/kg and 25mg/kg M102 groups when compared to vehicle dosed animals **(Fig. 3 B,C**; p <0.05 at 5mg/kg and <0.001 at 25mg/kg; one-way ANOVA followed by Dunnett’s multiple comparisons test).

The number of motor neurons in the lumbar ventral horns was counted and a significant dose-dependent increase in the number of surviving motor neurons was observed in the M102 groups when compared to vehicle treated animals (**Fig. 3D**; p <0.05 at 5mg/kg and <0.001 at 12.5 and 25mg/kg; one-way ANOVA followed by Dunnett’s multiple comparisons test). At 90 days of age there was a significant improvement in the CMAP amplitude of the M102 25 mg/kg dose group when compared to vehicle controls (**Fig. 3E, F** p-<0.05 one-way ANOVA followed by Dunnett’s multiple comparisons test). The percentage change in CMAP amplitude between 60 and 90 days of age showed a dose-dependent increase in the M102 dosed groups when compared to the vehicle dosed group with 1/7 mice showing an increase in in CMAP in the vehicle group versus 6/8 in the 25mg/kg M102 treated group (**Fig. 3G**).

### Dose prediction to humans and safety margin

Efficacy was observed in the mouse SOD1^G93A^ model at an oral dose of 12.5 mg/kg which resulted in a AUC_last_ of 419 ng*h/mL. Adjusting for the difference in human vs. mouse free fraction yields an estimated human equivalent exposure of 541 ng*h/mL. This value was used as the target efficacy exposure in the human dose projections as well as in the safety margin calculations. To confirm brain penetration of M102, additional plasma and brain PK studies were conducted in rats (**Suppl. Table 1**). After a single dose oral administration of 12 mg/kg M102, the brain-to-plasma ratios of M102 were determined to be 0.70 and 0.36 at 30 and 60 min post-dose, respectively. Subsequently, M102 was evaluated in non-Good Laboratory Practice (GLP) and GLP general toxicology studies in rats and non-human primates (NHPs). In the non-GLP toxicology studies, M102-related mild liver toxicity findings were observed at 250 mg/kg and 100 mg/kg in rats and NHPs, respectively. In 28-day GLP toxicology studies, 75 mg/kg was declared as the no observed adverse effect level (NOAEL) in both rats and NHPs. The free AUC on Day 28 at the NOAEL in both species, as well as the margin to the projected human free AUC necessary for efficacy, is 9.3-fold for rats, 10.1-fold for female NHPs, and 15.1-fold for male NHPs.

Human clearance, volume of distribution and half-life were estimated using PK data from rat and monkey studies. Various allometric scaling techniques were applied following the rule of exponents guidelines. Human clearance was also estimated from *in vitro* hepatocyte incubations. Correcting for brain weight, free fraction and using the multi-exponential method (*39*) resulted in estimates ranging from 55-111.7 mL/min/kg, while scaling intrinsic clearance from human hepatocytes resulted in an estimate of 84 mL/min/kg. Human volume of distribution was estimated by allometric scaling of preclinical species values and using an exponent of 1 and was approximately 81 L/kg. M102 has a bioavailability range of 10-30% in preclinical species. As a predictive model of human bioavailability does not exist, this range of potential values were modeled. Using the projected human efficacious exposure target of 302 ng*h/mL that was based on mouse efficacy at an oral dose of 12.5 mg/kg PO, a daily dose range in humans of 352-1417 mg is predicted.

### Increased oxidised RNA in affected CNS areas, and in the CSF and i-Astrocytes derived from ALS cases

Encouraged by the compelling data obtained in two *in vivo* models of ALS, we decided to evaluate the levels of oxidative stress detectable in patient biosamples and in a patient-derived cellular model of ALS, as potential biomarkers of target engagement and efficacy.

Oxidative stress is one of the hallmarks of ALS, with oxidised lipids and proteins being identified in post-mortem tissues (*40*). Due to the recently identified involvement of RNA dysregulation in ALS (*41*) and the known role for oxidized RNA in neurodegeneration (*42*), we aimed to map the presence of oxidized RNA in the areas of the CNS that are affected by the disease. As guanine is the base that is most susceptible to oxidation, we used an antibody against 8-oxo-2-oxyguanosine (8-OHG), as a marker of direct nucleic acid oxidation. Staining of frontal and motor cortex with adjacent white matter, as well as the cervical spinal cord, from 6 sALS cases, 4 cases carrying C9orf72 repeat expansion mutations and 3 healthy controls (case information in **Suppl. Table 2**) revealed that ALS cases display much higher levels of oxidised RNA compared to age-matched, neurologically unaffected controls (**Fig. 4 A,B**). Of note, cases carrying C9orf72 mutations with both ALS and fronto-temporal dementia (FTD), displayed significantly higher levels of oxidised RNA staining in the frontal cortex compared to both controls and ALS cases who displayed only a motor phenotype (**Fig. 4B**). Interestingly, although ALS cases displayed higher levels of oxidised RNA in the spinal cord compared to controls, amongst all the CNS areas analysed, the spinal cord was the tissue with the highest oxidative stress burden (**Fig. 4A**).

**Figure 4.**
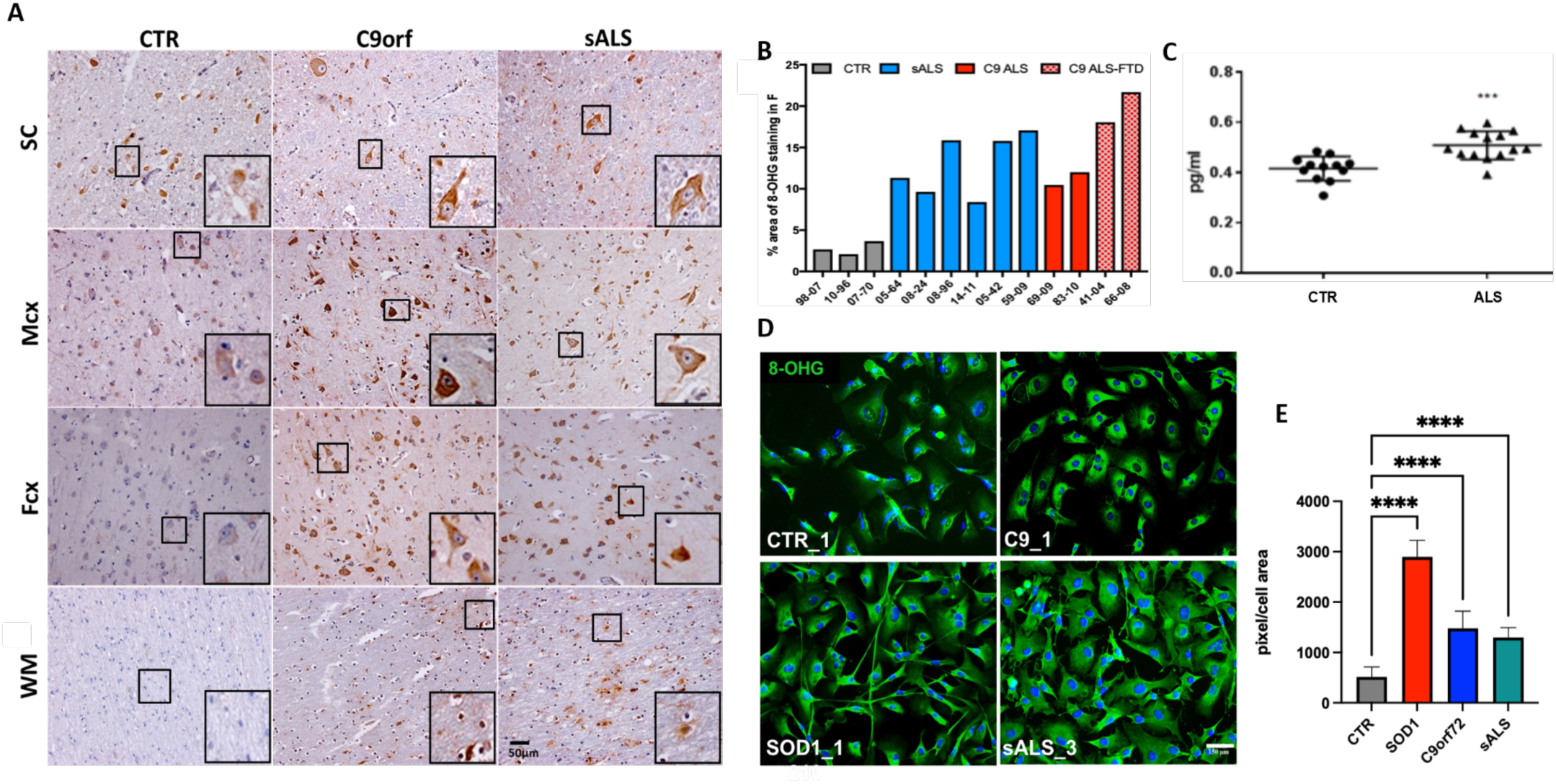
ALS cases display high levels of oxidized RNA compared to healthy controls. A) Representative images of 8-OHG staining in post-mortem tissue show increased oxidized RNA levels in C9-ALS and sALS patients compared to healthy controls, in several areas of the CNS. B) Respective quantifications in the frontal cortex (each bar represents one individual). C) Increased levels of oxidative stress are also detected in the CSF of ALS cases compared to healthy controls (N=14 ALS and N=12 controls). Unpaired t-test. D) Representative images of 8-OHG staining in iAstrocytes derived from healthy controls and SOD1-, C9- and sporadic ALS cases (scale bar: 150µm), and E) Respective quantifications show that patient-derived iAstrocytes also display high levels of oxidative stress, recapitulating what is observed in post-mortem tissue (N=2 controls; N=1 SOD1, N=3 C9-ALS; N=3 sALS). One-way ANOVA followed by Dunnett’s multiple comparisons test. SC: spinal cord; Mcx: motor cortex; Fcx: frontal cortex; WM: white matter. Significance: * <0.05; ** <0.01; *** <0.001, ****<0.0001.

Given the positive correlation between 8-OHG staining and pathology, we proceeded to examine whether oxidized RNA levels in the CSF could be used as a biomarker of oxidative stress. Using an ELISA kit specific for oxidized RNA, we detected higher levels of 8-OHG in the CSF of ALS cases (n=14; 10 males and 4 females; average age= 56.2 ± 10 years) compared to controls (n= 12; 5 males and 7 females; average age= 44.2 ± 12 years (**Fig. 4C**, p<0.01 unpaired t-test). However, there did not seem to be any correlation between clinical/genetic characteristics and levels of oxidized RNA (subject information in **Suppl. Table 3**).

To determine whether increased levels of oxidized RNA were also present in ALS patient-derived astrocytes, we stained iAstrocytes differentiated from induced neural progenitor cells (iNPCs) directly reprogrammed from fibroblasts collected via skin biopsy from ALS cases and age-matched healthy controls (subject information in **Suppl. Table 4**). These iAstrocytes are known to retain the features of ageing that are likely to contribute to patient-specific phenotypes (*43*), and therefore constitute a robust *in vitro* model of the astrocyte contribution to ALS. iAstrocytes derived from SOD1, C9orf72 and sporadic ALS patients displayed higher levels of oxidized RNA compared to healthy controls (**Fig. 4D,E**, p<0.001 one-way ANOVA followed by Dunnett’s multiple comparisons test), recapitulating the findings in post-mortem tissue.

### M102 treatment is a dual activator of the NRF2 and HSF1 pathways in ALS patient-derived iAstrocytes, and is sufficient to reduce oxidative stress, misfolded SOD1, and TDP43 proteinopathy in vitro

As ALS patient-derived iAstrocytes recapitulate the high levels of oxidative stress observed in post-mortem tissue, we next set out to evaluate the ability of M102 treatment to activate the antioxidant NRF2 pathway and therefore contribute to decreased oxidative stress in these cells.

It is known that the NRF2 pathway is downregulated in astrocytes from models of ALS (*27*). Consistently, iAstrocytes derived from SOD1 (n=1), C9orf72 (n=3) and sporadic (n=2) ALS cases displayed lower levels of NQO1, a downstream target of NRF2 (*18, 20*), compared to age-matched healthy controls, under baseline conditions (**Fig. 5A,B**, p <0.05 one-way ANOVA followed by Dunnett’s multiple comparisons test). Importantly, when treated with 10 μM of M102 for 48h, the levels of NQO1 increase significantly in iAstrocytes from healthy controls and ALS cases, including SOD1, C9orf2 and sporadic patient lines (**Fig. 5A,C**, p <0.01 two-way ANOVA followed by Šídák’s multiple comparisons test). Interestingly, iAstrocytes from patients with a C9orf72 mutation showed a higher response to M102 compared to iAstrocytes from sporadic cases or cases with a SOD1 mutation. Overall NRF2 levels in the nucleus vs cytoplasm also increased upon 24h treatment with M102 (**Figure 5D, E**, treatment effect p< 0.01 two-way ANOVA), providing evidence that M102 activates the NRF2-ARE pathway in ALS patient-derived iAstrocytes. Similar to what is observed in mouse models of ALS, exposure to M102 also leads to increased expression of HSF-1 in patient-derived iAstrocytes **(Fig. 5F, G**, treatment effect p <0.001 two-way ANOVA), indicating that M102 is also a dual activator of the NRF2-ARE and HFS-1-HSE pathways *in vitro*.

**Figure 5.**
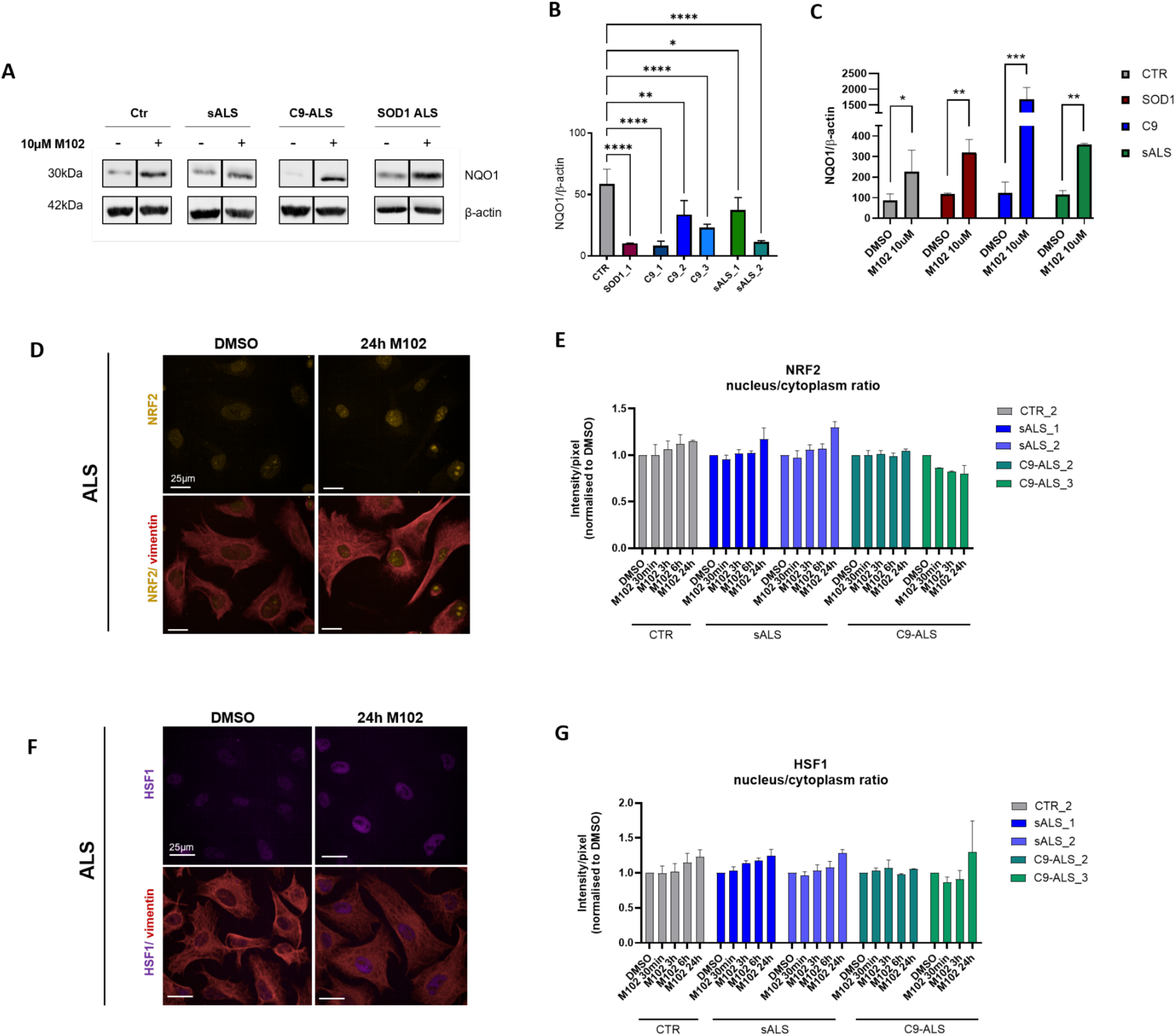
M102 treatment also activates both NRF2 and HSF1 pathways in sporadic, C9, and SOD1 patient-derived iAstrocytes. A) Representative western blots showing increased expression of NQO1 upon 10 μM M102 treatment for 48h in sporadic, C9 and SOD1 ALS patient-derived iAstrocytes. DMSO and M102 treated samples of each line are from the same blot but were cropped because there were different conditions in between these lanes. Original full blots are available upon request. B) Quantification shows that iAstrocytes derived from ALS cases with different genotypes display lower baseline levels of NQO1 compared to healthy controls (N=3 technical repeats per cell line; one-way ANOVA followed by Dunnett’s multiple comparisons test), and C) These levels are increased upon 10 μM M102 treatment for 48h (N=3 technical repeats per cell line; two-way ANOVA followed by Šídák’s multiple comparisons test). (D) Representative staining of NRF2 before and after 10 μM M102 exposure in ALS iAstrocytes, and respective quantifications under (E) Baseline levels and in response to M102 treatment for 30min to 24h (N=4-5 technical repeats per cell line). Two-way ANOVA treatment effect: p-value=0.0067. (F) Representative staining of HSF1 before and after 10 μM M102 treatment in sALS and C9-ALS iAstrocytes, and respective quantifications under (G) Baseline levels and upon M102 treatment for 30min - 24h (N=4-5 technical repeats per cell line). Two-way ANOVA treatment effect: p-value=0.0003. Significance: * <0.05; ** <0.01; *** <0.001.

Consistent with these findings, iAstrocytes derived from SOD1, C9orf72 and sporadic ALS cases treated with 10 μM M102 for 48h showed decreased levels of oxidized RNA, thus confirming that M102 is capable of reducing the levels of oxidative stress in ALS astrocytes (**Fig. 6A,B**, p<0.01 two-way ANOVA followed by Šídák’s multiple comparisons test).

**Figure 6.**
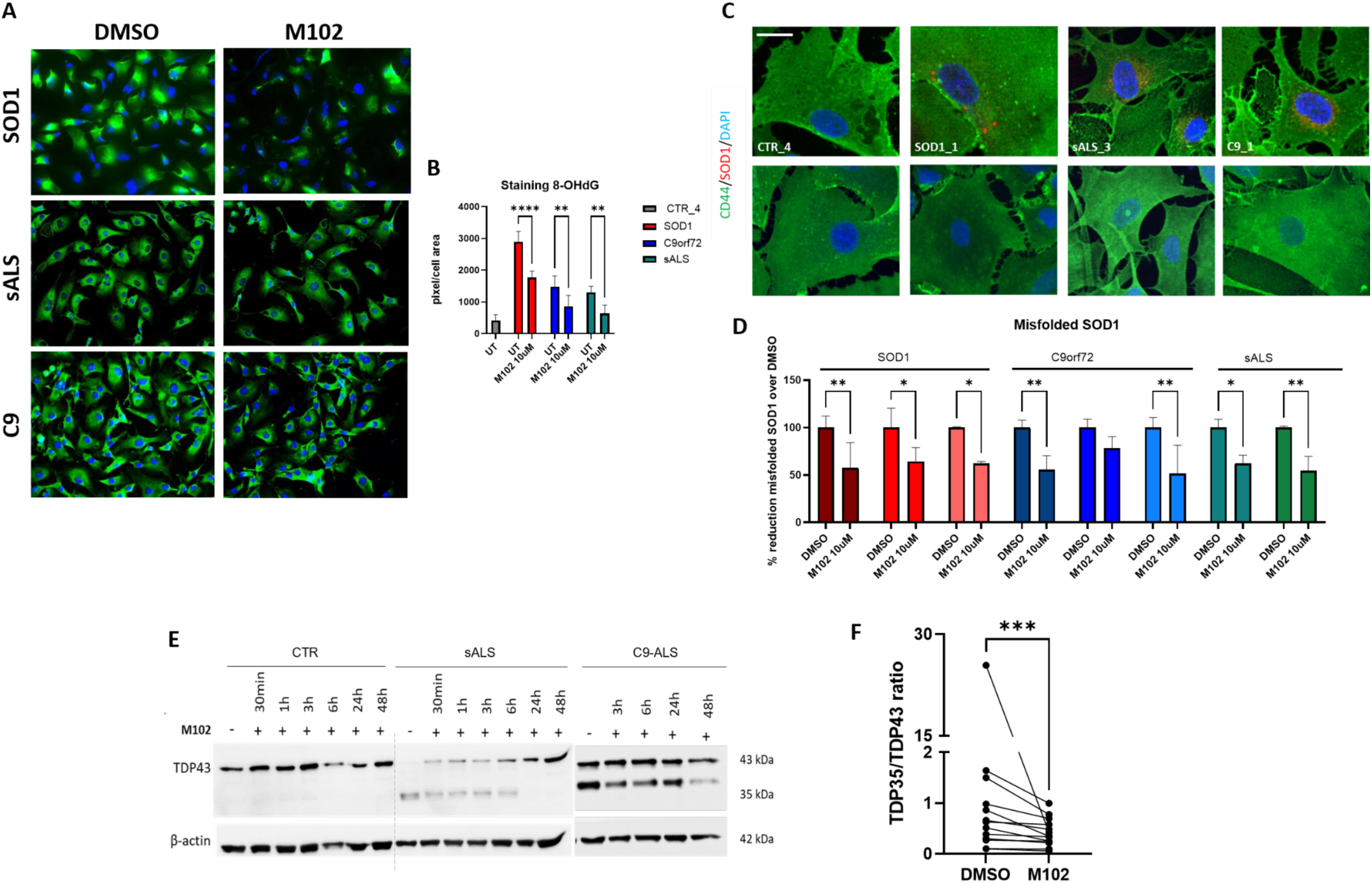
M102 treatment leads to reduction of oxidized RNA, misfolded SOD1 and TDP-43 proteinopathy in ALS patient-derived iAstrocytes. A) Representative images of 8-OHdG staining and (B) Respective quantifications show reduced levels of oxidized RNA upon 10 μM M102 exposure for 48h in iAstrocytes derived from SOD1, sporadic and C9 ALS patients. N=2-3 per genotype. Two-way ANOVA followed by Šídák’s multiple comparisons test. C) Representative images of misfolded SOD1 before (top panel) and after (bottom panel) exposure to M102 and D) Respective quantifications show reduction of misfolded SOD1 in iAstrocytes derived from SOD1, sporadic and C9 ALS patients upon M102 exposure (N=3 SOD1 ALS, N=3 C9-ALS, N=2 sALS - each with 3 technical repeats). Two-way ANOVA followed by Šídák’s multiple comparisons test. E) Representative immunoblotting images and F) Respective quantifications show a time-dependent reduction of TDP-43 proteinopathy upon M102 treatment (10 μM for 48h) in sALS (n=10, 2-4 technical repeats each) and C9-ALS (n=3, 3-4 technical repeats each; paired t-test) patient-derived iAstrocytes, characterised by a reduction of the fragmented 35kDa band. Significance: * <0.05; ** <0.01; *** <0.001.

The HSF-1-HSE pathway is known to play an essential role in maintaining proteostasis by facilitating protein folding and avoiding protein misfolding (*35*). As we showed that M102 is a strong activator of the HSF-1 pathway, we set out to investigate whether M102 treatment has a beneficial effect on the reduction of misfolded SOD1 and TDP-43 proteinopathy, two hallmarks of ALS pathology (*44, 45*). By using an antibody raised against misfolded SOD1 (B8H10), we detected perinuclear staining in iAstrocytes from SOD1 cases, as well as cases carrying C9orf72 mutations and sALS cases, consistent with previous findings in post-mortem CNS tissue. Upon treatment with M102, we observed a significant reduction of misfolded SOD1 in iAstrocytes derived from SOD1, C9orf72, and sporadic ALS cases (**Figure 6C,D**, p <0.05 in 8 out of 9 cell lines; two-way ANOVA followed by Šídák’s multiple comparisons test). Induced astrocytes recapitulate one of the key hallmarks of ALS, i.e. TDP-43 proteinopathy, detected as the presence of TDP-43 fragments (observed at 35 kDa) in sporadic and C9orf72 ALS patient iAstrocytes (**Fig. 6E**), but not in SOD1 cases (data not shown). Importantly, we observe that time-dependent treatment with M102 leads to a reduction of TDP-43 proteinopathy, especially following 48h exposure (**Fig. 6E, F**, p <0.01 paired t-test).

### M102 rescues MN survival in co-culture with ALS patient-derived iAstrocytes by targeting multiple mechanisms known to underlie ALS pathophysiology

Astrocytes from ALS cases are known to be toxic to MNs, contributing to death of healthy motor neurons in co-culture (*10*). We have previously reported that iAstrocytes differentiated from iNPCs directly reprogrammed from fibroblasts of SOD1, C9orf72 and sporadic ALS cases are toxic to MNs (*10*). As M102 seems to target multiple mechanisms associated with ALS, including oxidative stress and protein misfolding and aggregation, we asked whether M102 treatment would be sufficient to rescue MN survival when in co-culture with toxic ALS astrocytes. For this, we used our previously described iAstrocyte-MN co-culture model (*10*). Briefly, ALS patient-derived iAstrocytes were treated with DMSO or M102, using the Echo 550 liquid dispenser, and 24h later co-cultured with healthy mouse MNs expressing GFP under a Hb9 promoter. The MNs were then scanned in an InCell Analyser 24h and 72h after treatment, and the numbers of viable MNs counted (**Fig. 7A**).

**Figure 7.**
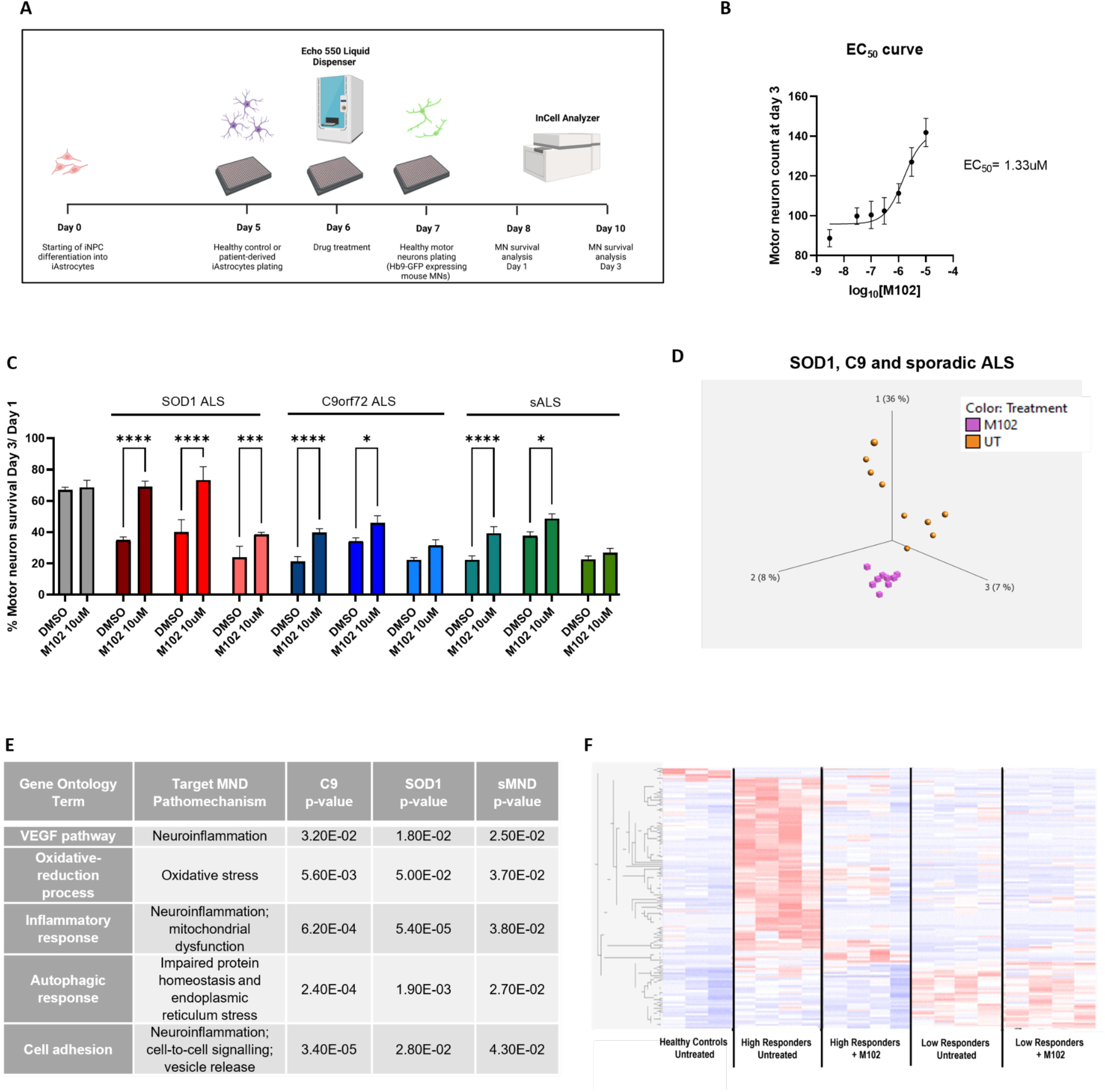
M102 treatment rescues MN survival in co-cultures with C9, SOD1 and sporadic ALS patient-derived astrocytes, and targets multiple pathways known to be associated with ALS. A) Schematic figure of the MN-iAstrocyte co-culture system previously described (*70*). B) EC50 calculated from dose-response curve tested in sALS patient-derived iAstrocyte/MN co-cultures (average of 4 cell lines, 6-8 technical repeats each). C) ALS-patient derived iAstrocytes are toxic to co-cultured MNs under basal conditions, leading to increased MN death compared to healthy controls, and motor neuron survival is rescued upon 10 μM M102 treatment for 48h in sporadic, C9 and SOD1 ALS cases. N=3. (Two-way ANOVA followed by Šídák’s multiple comparisons test). D) RNA sequencing from iAstrocytes treated with 10 μM M102 or DMSO for 48h reveal transcriptomic changes in response to M102 treatment in ALS iAstrocytes. E) Enriched pathways in response to M102 exposure identified in common between C9, SOD1 and sALS iAstrocytes show that M102 treatment targets various pathophysiological mechanisms known to play a role in ALS. F) Heatmap shows the transcriptomic profile of high and low responders (i.e. >50% rescue and <50% rescue, respectively) to M102 under baseline conditions and upon treatment with 10 μM M102 for 48h. Significance: * <0.05; ** <0.01; *** <0.001.

To assess the optimal dose of M102 to use, we started by testing 6 different concentrations of M102, ranging from 0.03 μM to 10 μM, in 4 sALS iAstrocyte lines. As expected, under baseline conditions, MNs co-cultured with ALS iAstrocytes showed reduced survival compared to MNs co-cultured with iAstrocytes derived from healthy controls. Furthermore, our dose-curve response showed that M102 has an EC_50_ of 1.33 μM and the maximum neuroprotective effect in co-culture was achieved with a dose of 10 μM (**Fig. 7B**). We therefore used 10 μM M102 to further evaluate whether M102 is sufficient to rescue MN survival when in co-culture with multiple iAstrocyte lines derived from both familial and sporadic ALS cases. Under baseline conditions, MNs co-cultured with ALS patient-derived iAstrocytes displayed reduced survival (between 30%-60% reduction in MN survival depending on the patient donor) compared to MNs co-cultured with iAstrocytes derived from healthy controls (**Fig. 7C**). Treatment with 10 μM M102 led to a significant increase in MN survival in co-cultures with 7 out of 9 different iAstrocyte patient lines, including SOD1, C90rf72 and sporadic cases (**Fig. 7C**, p <0.05 two-way ANOVA followed by Šídák’s multiple comparisons test). Interestingly, the response to M102 varied between patient lines, with 4 patient lines responding more strongly by increasing MN survival by >50% compared to baseline levels

In order to further explore the neuroprotective mechanisms of M102 and better understand how M102 could play a role in ameliorating iAstrocyte toxicity and MN survival, we performed RNA Sequencing in patient-derived iAstrocytes before and after 10 μM M102 exposure for 48h. Our data show that there is a clear separation of the transcriptomic profile before and after M102 treatment, as shown by a principal component analysis (PCA) plot (**Fig. 7D**). This transcriptomic shift in response to M102 is observed in iAstrocytes derived from SOD1, C9orf72, and sporadic cases (**Suppl. Fig. 4 A,B,C**, respectively). Differential expression analysis between treated and untreated iAstrocytes identified a total of 160, 283, and 267 differentially expressed genes (DEGs) in C9-ALS cases, SOD1 cases, and sALS cases, respectively (**Suppl. Fig. 4D**). We further performed gene ontology analysis to investigate the predominant pathways affected by M102 treatment. The results showed that M102 treatment altered the expression of genes associated with neuroinflammation, mitochondrial dysfunction, autophagic response and cell adhesion (**Fig. 7E, Suppl. Table 5**), all known to be important drivers of the pathophysiology of ALS (*9*). Taken together with our results demonstrating that M102 is a dual activator of the NRF2 and HSF-1 pathways, these results indicate that M102 targets multiple pathophysiological mechanisms of ALS.

As we observed that some iAstrocyte lines responded more strongly to M102 than others, we next set out to evaluate whether we could discriminate high and low responders to M102 based on the individual patient transcriptomic profiles. For this, we classified the iAstrocytes in ‘high’ and ‘low’ responders to M102 based on the levels of motor neuron survival upon M102 treatment in iAstrocyte-MN co-cultures. iAstrocytes that responded to M102 with an increase in motor neuron survival of over 50% were considered ‘high responders’ (4 patients) whereas the iAstrocytes that responded with an increase in motor neuron survival of less than 50% were considered ‘low responders’ (4 patients). A heatmap of healthy controls and ALS patient-derived iAstrocytes before and after M102 treatment showed that a set of 161 transcripts identified significant differences between “high” and “low” responders to M102. Interestingly, the low responders to M102 show a transcriptomic profile for these 161 transcripts that is closer to healthy controls compared to high responders, under baseline conditions. Upon M102 treatment, high responders displayed a shift in their transcriptomic profile for these 161 transcripts, which became more similar to the profile of healthy controls (**Fig. 7F**), thus indicating that a subgroup of these genes could be used as a biomarker of drug response.

## MATERIALS AND METHODS (For Full Materials and Methods, see Supplementary Material)

### Study design

This study aims to provide a comprehensive package of preclinical efficacy data for M102, a CNS penetrant small molecule electrophile capable of activating both NRF2-ARE and HSF-1-HSE pathways, *in vivo* and *in vitro*. To do this, the pharmacokinetic profile was assessed in C57Bl/6 mice; GLP general toxicology studies were performed in rats and NHPs; and target engagement and efficacy were assessed across two ALS mouse models: TDP-43^Q331K^ and SOD1^G93A^ dosed with M102 subcutaneously and orally respectively. RT-qPCR was used to assess target engagement via gene expression changes after M102 administration. Immunohistochemistry (IHC) was used to identify and count the number of motor neurons in the spinal ventral horn after M102 treatment. Compound muscle action potential (CMAP) amplitude of hind limb muscles, rotarod performance, and gait parameters were used to assess M102 efficacy on readouts of motor function. Together, these data enabled prediction of human efficacious exposures and doses, which were established to be well within the safety margin predicted from GLP toxicology studies.

Post-mortem tissue, human CSF samples, and ALS patient-derived astrocytes were used to confirm that ALS cases present higher levels of oxidative stress compared to healthy controls, and that M102 is capable of reducing the levels of oxidative stress. A combination of immunocytochemistry (ICC) and western blotting analysis was used to assess the effects of M102 on NRF2-ARE and HSF-1-HSE pathway activation, as well as to assess TDP43 proteinopathy in patient-derived astrocytes. MN-astrocyte co-cultures were performed to assess whether M102 can rescue motor neuron survival in the presence of toxic ALS iAstrocytes, and to determine high and low responders to M102. RNA Sequencing of untreated and M102 treated patient-derived iAstrocytes was used to identify additional biological mechanisms targeted by M102.

No study size calculations or randomization were carried out. All samples were quantified in a blinded manner for microscopy and mouse experiments. No cell or animal samples were excluded.

### Ethics statement

GLP general toxicology studies in rats and NHPs to evaluate potential toxicity of M102 were conducted at WuXi AppTec (Suzhou, China). The protocol and any amendments or procedures involving the care or use of animals in this study were reviewed and approved by the Institutional Animal Care and Use Committee (IACUC) prior to the initiation of such procedures. A staff veterinarian monitored the study for animal welfare issues.

All mouse studies were carried out under a UK Home Office project license by individuals that held the appropriate UK Home Office personal license and had appropriate training for procedures. All work was carried out under the terms of the UK Animals (Scientific Procedures) Act 1986 and animals were housed and maintained in line with Home Office Code of Practice for House and Care of Animals Used in Scientific Procedures.

Formalin-fixed, paraffin-embedded (FFPE) human CNS post-mortem tissue was obtained from the Sheffield Brain Tissue Bank with Research Ethics Committee approval (Sheffield Brain Bank –SBB-, Ethics Committee reference 08/MRE00/103). Human CSF was obtained from the University of Sheffield Biobank with Research Ethics Committee approval number STH16573. Fibroblasts were collected from skin biopsies donated by ALS patients and controls with informed consent (Ethical Committee approval references: 12/YH/0330; 16/LO/2136).

### Mice

Mice were housed in same sex groups of between 2 and 5 mice per cage. Each cage consisted of a plastic house, sawdust covering the floor (Datesand) and paper wool bedding (Datesand). The mice had *ad libitum* access to water and food (standard rodent diet 2018, Envigo). Temperatures in the rooms were maintained at 21°C with a 12 hour light/dark cycle (7am – 7pm). Wild type animals used were C67BL/6 mice either from Envigo or non transgenic mice from the SOD1^G93A^ colony.

The SOD1^G93A^ C57BL/6 transgenic mice were bred in-house. The B6SJL-Tg(SOD1-G93A)1Gur/J mice were backcrossed onto the C57BL/6J OlaHsd background for at least 20 generations. This line has been extensively characterised in-house and develop a reproducible progressive motor phenotype (*46*). The TDP-43^Q331K^ C57BL/6NJ transgenic mice were bred in-house and were obtained from Jackson laboratory (stock number 017933). These mice were originally on a C57BL/6NCrl background (*47*) but have been crossed onto a C57BL/6NJ background for a minimum of 4 generations and have been extensively characterised by the authors (*32*). Mice were ear-clipped for identification and genotyping. Genotyping for the two colonies was carried out as previously described (*32, 46*).

### Differentiation of patient-derived neurons, motor neurons and astrocytes

#### Tissue culture to generate induced neural progenitor cells (iNPCs) and iAstrocytes from donated fibroblasts

Fibroblasts collected from skin biopsy material donated by ALS cases and controls were directly reprogrammed into iNPCs, using a combination of retroviral vectors (Oct3/4, Sox2, Klf4, and c-Myc), as previously described (*10*). Briefly, the fibroblasts were treated with 700 μL medium/viral vector overnight, then washed 2× with PBS, and fed once per day with fibroblast medium (DMEM plus 10% FBS) for 3 days. At day 4 the cells were switched to NPC conversion medium consisting of DMEM/F12, 1% N2, 1% B27, 20 ng/mL FGF2, 20 ng/mL EGF, and heparin (5 μg/mL; Sigma-Aldrich) and fed every day thereafter. When the cells changed shape and presented sphere-like structures, they were lifted with accutase, centrifuged, resuspended in NPC conversion medium, and replated for expansion. Once the NPC culture was established, the medium was switched to NPC medium consisting of DMEM/F12, 1% N2, 1% B27, and FGF2 (40 ng/mL). To differentiate iNPCs into iAstrocytes, iNPCs were seeded in NPC medium at low density in a fibronectin-coated 10-cm dish. The day after, the medium was changed to DMEM containing 10% FBS and 0.3% N2 and the cells were allowed to mature for at least 7 days.

### Motor neuron differentiation from embryonic stem cells

Mouse embryonic stem cells expressing GFP under the MN-specific promoter HB9 (HBG3 cells; kind gift from Tom Jessell, Columbia University, New York) were cultured on primary mouse embryonic fibroblasts (Millipore). For differentiation into MNs, cells were lifted with trypsin and resuspended in DFK10 culture medium consisting of knockout DMEM/F12, 10% knockout serum replacement, 1% N2, 0.5% L-glutamine, 0.5% glucose (30% in water), and 0.0016% 2-mercaptoethanol. The cells were plated on nonadherent Petri dishes to allow formation of embryoid bodies. After 1 d of recovery, 2 μM retinoic acid (Sigma) and 1 μM smoothened antagonist (SAG, Merk) were added every day with fresh medium. After 5 d of differentiation, the embryoid bodies were dissociated and sorted for GFP on a BD FACSVantage/DiVa sorter.

### Human iAstrocyte-murine motor neuron co-culture assay

Human plasma fibronectin (Merck Millipore) was diluted 1:400 in PBS, and 5 µL was added per well on 384-well plates (Greiner Bio-one, 781091. Plates were coated for at least 5 min at RT. A total of 2,000 human iAstrocytes were seeded in 35 μL iAstrocyte media per well on fibronectin-coated 384-well plates. Plates were centrifuged at 1,760 x g for 60 s, and cells were incubated for 24 h. Drug was then delivered to iAstrocytes in 100 % anhydrous DMSO (Sigma, 276855) using an Echo550 liquid handler (Labcyte). The final concentration of DMSO was 0.1 % (v/v) in the media in all wells. Plates were centrifuged at 1,760 x g for 60 s, and cells were incubated for a further 24 h. A total of 2,500 murine Hb9-GFP+ motor neurons were seeded per well in motor neuron media (KnockOut DMEM (45% v/v), F12 medium (45% v/v), KO Serum Replacement (10% v/v), 50 units/ml penicillin/streptomycin (Lonza), 1 mM L-glutamine, 1X N-2 supplement (Thermo-Fisher Scientific), 0.15% filtered glucose, 0.0008% (v/v) 2-mercaptoethanol, 20 ng/ml GDNF, 20 ng/ml BDNF, 20 ng/ml CNTF) on top of the pre-treated iAstrocytes. Plates were centrifuged at 1,760 x g for 60 s. Hb9-GFP+ motor neurons were imaged after 24 and 72 hours using an INCELL analyser 2000 (GE Healthcare), and the number of viable motor neurons was counted using the Columbus™ analysis software (Perkin Elmer). The number of viable motor neurons (defined as GFP+ motor neurons with at least 1 process) that survived after 72 hours in co-culture was calculated as a percentage of the number of viable motor neurons after 24 hours in co-culture. The percentage survival of motor neurons was then normalised to the DMSO control for each individual iAstrocyte line.

### Statistical analysis

Statistical analysis was performed using GraphPad Prism v11.00. Normality was assessed by the Shapiro-Wilk test. Unpaired t-test or the Mann Whitney test were used to compare 2 data points, depending on whether the samples presented a Gaussian distribution or not, respectively. Paired t-test was used to compare the same samples before and after treatment. One-way ANOVA followed by Dunnett’s multiple comparisons tests was used to compare more than 2 data points in 1 group. Two-way ANOVA followed by Šídák’s multiple comparisons tests was used to compare more than 2 data points in 2 groups.

## DISCUSSION

Our understanding of the genetic underpinnings and pathophysiological mechanisms in ALS has substantially increased in recent years (*9*). However, apart from the recent promising emergence of tofersen as a disease modifying therapy for the two percent of ALS patients who harbor mutations in the *SOD1* gene, other approved drugs have only marginal effects on life expectancy (riluzole) or indices of disease progression (edaravone) (*48*). Therapies targeting individual pathways have failed to generate significant benefit during several decades of clinical trials. Previous attempts at identifying effective neuroprotective therapies for ALS have failed, as they do not account for disease heterogeneity and the multiple mechanisms driving motor neuron injury (*17, 49*). Data from disease model systems and from human biosamples provide strong evidence for a role of redox imbalance (*40*), inflammation (*50*), mitochondrial dysfunction (*51*) and altered proteostasis, including autophagy and mitophagy (*51, 52*), as four key drivers in the pathobiology of ALS (*9, 49*). Few, if any, studies in ALS have shown multi-target engagement for the proposed therapeutic agent in the clinic (*17*).

We previously identified M102, a CNS-penetrant, small molecule, activator of the NRF2-ARE pathway. Here, we have also shown that it also activates the HSF-1-HSE transcription factor pathway, which targets multiple genetically validated pathophysiological mechanisms in ALS. We identified M102 in a screen to discover CNS penetrant NRF2-ARE pathway activators (*28*). M102 (S-apomorphine hydrochloride hemihydrate) is a proprietary new chemical entity (NCE) and the S-enantiomer of the marketed R-apomorphine (Apokyn®; pure R-enantiomer). The R-enantiomer is a dopamine agonist administered subcutaneously for the management of advanced Parkinson’s disease. M102 is a very weak dopamine antagonist and does not show the adverse effects associated with dopamine agonism (*29*). Initial work showed beneficial effects in the SOD1^G93A^ mouse model and in indices of oxidative stress in ALS patient fibroblasts (*28*). Here, we demonstrate that M102 is a dual activator of NRF2 and HSF1 transcription factor pathways, two upstream master regulators of neuroprotective mechanisms, with the potential to modulate all four of these key drivers of neurodegeneration and with excellent penetration across the blood brain barrier (*34, 39*).

NRF2 is a stress-responsive transcription factor and a master regulator of anti-oxidant and anti-inflammatory genes, activating the transcription of >1000 cytoprotective genes and with a multi-modal contribution to healthy mitochondrial function and regulation of key autophagy genes (*53, 54*). These pleiotropic effects enable the cell to maintain homeostasis under stress conditions. NRF2 activity is tightly regulated via a complex set of transcriptional and post-translational controls, principally by KEAP1 in the cytoplasm and BACH1 in the nucleus (*55*). Under physiological conditions, KEAP1 promotes constitutive proteasomal degradation of Nrf2 (*56*) while BACH1 acts as a transcriptional antagonist of NRF2 target genes (*57*). Under oxidative stress conditions, KEAP1 is inactivated by modification of its reactive cysteine residues allowing Nrf2 to escape degradation and newly synthesized Nrf2 is able to translocate to the nucleus and bind to the anti-oxidant response elements (AREs) of multiple cytoprotective genes. This cytoprotective gene expression response provides protection against several key pathophysiological mechanisms operating in ALS including oxidative stress, mitochondrial dysfunction, inflammation and dysregulated proteostasis (*18*). The stress response of the KEAP1-NRF2-ARE system is stronger in astrocytes compared to neurons (*58*). A body of evidence from *in vitro* and *in vivo* model systems and from post-mortem CNS tissue from ALS patients has indicated that the NRF2 response is impaired in ALS and has also been shown to decline with age (*59–61*).

HSF-1 is a stress-inducible transcription factor that is the key driver for the expression of multiple heat shock proteins which act as chaperones responsible for the correct folding of newly synthesized proteins, the refolding of denatured proteins and the prevention of aggregation of misfolded proteins. This heat shock response is a pro-survival pathway activated under conditions of cellular stress. This cellular stress response declines with age, with decreased ability of HSF-1 to bind to the promotors of genes encoding heat shock proteins (*35*). Motor neurons have been reported to display a high threshold for the induction of the heat shock stress response (*62*). Nevertheless, activation of the HSF- 1/HSP pathways has shown beneficial effects in cellular and animal models of ALS, including clearance of TDP-43 aggregates (*63*). Activation of HSF-1 is an attractive pharmacological target for ALS and other neurodegenerative conditions characterized by proteotoxic stress. However, to date, many small molecule activators of HSF-1 have shown undesirable properties *e.g*. by acting as Hsp90 inhibitors or by exerting direct proteotoxic effects (*35, 64*).

*In vitro* and *in vivo* data presented here show that M102 is a multi-target drug with predicted neuroprotective benefits for the treatment of ALS. As a dual activator of NRF2 and HSF1 pathways, M102 is able to activate anti-oxidant and anti-inflammatory pathways, as well as beneficial modulating effects on proteostasis and mitochondrial function. *In vitro* data also show that M102 can reduce TDP-43 proteinopathy and biomarkers of oxidative stress in patient-derived astrocytes. Moreover, M102 is capable of rescuing MN survival in co-cultures with iAstrocytes derived from *C9orf72*, *SOD1*, and sporadic ALS patients, as well as spinal cord MNs in the SOD1^G93A^ mouse model. The efficacy of M102 was shown *in vivo* in both *SOD1*^G93A^ and *TDP-43*^Q331K^transgenic mouse models of ALS, with beneficial effects on a translatable neurophysiological biomarker (CMAP) which correlates with motor neuron survival in the spinal cord. The transcriptome of patient-derived astrocytes exposed *in vitro* to M102 confirm the activation of multiple key neuroprotective pathways in astrocyte lines from several subgroups of ALS. There is a degree of heterogeneity in the level of the *in vitro* response to M102, with the transcriptome of high responders shifting substantially towards the profile observed in the astrocytes of healthy controls.

This comprehensive package of pre-clinical efficacy data for therapy development in ALS is accompanied by strong safety data. We have completed the process of Good Manufacturing Practice (GMP) manufacturing for M102 and a series of GLP toxicology and safety pharmacology studies in rats and non-human primates, which have demonstrated that M102 will have an acceptable safety margin (9.3-15.1x) in clinical studies. Pharmacokinetic studies show strong penetration of M102 across the blood-brain-barrier, with a brain-plasma ratio of 0.7 at 30 min post-dosing.

Other NRF2 activators have been investigated in clinical trials or have been approved for medical use. These include dimethylfumarate (DMF) (Tecfidera®, Biogen) and omaveloxolone (Reata). DMF was originally approved for the treatment of psoriasis (Fumaderm^®^) and was later repurposed for the treatment of relapsing-remitting multiple sclerosis (Tecfidera^®^). A phase 2 trial of DMF in ALS provided Class 1 evidence of safety at a dose of 480mg/day and lack of disease-modifying efficacy (*65*). DMF treatment is associated with dose-limiting lymphopenia and flushing (Tecfidera® Prescribing Information). Omaveloxolone (Skyclarys^®^) is a potent NRF2 activator that has been approved by the FDA for the treatment of Friedreich’s ataxia. By activating the NRF2 pathway, omaveloxolone ameliorates oxidative stress and improves mitochondrial function. As a potent NRF2 activator, omaveloxolone exhibited significant liver toxicity with elevated AST/ALT levels in 37% of patients exposed to a dose of 150 mg (*66*). Toxicity has also been reported with other potent NRF2 activators, such as bardoxolone methyl (EC_50_: 53 nM) which showed significant heart, liver, and renal toxicity in humans (*67*). In contrast, our preclinical toxicological studies indicate that M102 has a much higher safety margin in relation to liver toxicity. Arimoclomol is a co-activator of HSF-1 and has been ineffective in two clinical trials targeting *SOD1*-ALS initially and then ALS more broadly (*68,69*).

There is a huge unmet need for more effective neuroprotective therapies to slow disease progression in ALS. Here, we have demonstrated that M102, a combined activator of NRF2 and HSF1 signaling pathways, has positive therapeutic effects in two different ALS transgenic mouse models and improves motor neuron survival and multiple pathological markers (i.e. oxidative stress, misfolded SOD1, TDP43 proteinopathy) in a range of human ALS cellular model systems. Importantly, these neuroprotective effects are seen across multiple subtypes of ALS, including the two most common subtypes caused by mutations in the *C9orf72* and *SOD1* genes, as well as sporadic ALS cases. Taken together, we have demonstrated that M102 has the potential to modulate multiple key drivers of neurodegeneration, increasing the chances of achieving impactful neuroprotection and disease modifying effects in ALS. Its positive effect on TDP-43 proteinopathy and on rescuing motor neuron survival both *in vivo* and in co-culture with iAstrocytes derived from both familial and sporadic ALS patients, also suggest that M102 may be beneficial for a wide range of ALS subtypes as ∼97% of ALS cases display TDP-43 proteinopathy and ∼90% of cases are sporadic.

## General

We are very grateful to the ALS patients and healthy control subjects who generously donated biosamples to support this work.

## Funding

Medical Research Council Developmental Pathway Funding Scheme: MR/V027735/1 (RJM, PJS, LF, NS) FightMND Australia grant: 03_DDG_2020_Shan (NS, PJS, RJM, LF).

Amyotrophic Lateral Sclerosis Research Program supported by the Assistant Secretary of Defense for Health Affairs endorsed by the Department of Defense: W81XWH2210175 (NS, PJS, RJM, LF).

Motor Neuron Disease Association: A Multi-Centre Biomarker Resource Strategy in ALS (AMBRoSIA). MNDA 972-797(PJS).

NIHR Sheffield Biomedical Research Centre: NIHR 203321 (PJS).

## Author contributions

Conceptualization: PJS, RJM, LF, NS.

Methodology: AFK, RRM, CFA, KB, SM, TM, MM, NT, SNB, AS, SS, SNM, AD, TW, RJM, LF, PJS.

Investigation: AFK, RRM, CFA, KB, SM, TM, MM, NT, SNB, AS, SS, SNM, AD, TW, RJM, LF, PJS.

Visualization: AFK, RRM, CFA, KB, SM, TM, MM, NT, SNB, AS, SS, SNM, AD, TW, RJM, LF, PJS.

Funding acquisition: PJS, RJM, LF, NS.

Project administration: PJS, RJM, LF, NS.

Supervision: PJS, RJM, LF, NS.

Writing – original draft: PJS, RJM, LF, NS, AFK, RRM, CFA

Writing – review & editing: All authors

## Competing interests

PJS is a member of the Scientific Advisory Board for Aclipse Therapeutics and PJS, RJM and LF are share-holders of Aclipse Therapeutics.

NS and INK are employees of Aclipse Therapeutics.

### Patents relevant to M102

#### PCT/US2019/056996 - Treatment of Neurodegenerative diseases

Inventors: Ning Shan, Richard Mead, Laura Ferraiuolo, Pamela J Shaw. This covers the mechanism of action of a drug identified at SITraN for the treatment of neurodegenerative diseases, including ALS. Submitted in 2019.

#### PCT/US2019/056998

Treatment of Neurodegenerative diseases. Inventors: Laura Ferraiuolo, Ning Shan, Pamela J Shaw. This covers the specific properties of a neuroprotective compound identified at SITraN. Submitted in 2019.

PCT/US2020/45321 - Pharmaceutical Composition For Use In The Treatment Of Neurological Diseases. Inventor: Ning Shan. This covers the pharmaceutical compositions and associated pharmacokinetics of a drug identified at SITraN. Submitted in 2019.

## Data and materials availability

Materials used in this study can be made available subject to Materials Transfer agreements (MTAs). RNA Sequencing data will be made available on a publically available database following publication of the manuscript. All other data are available in the manuscript main text and supplementary materials.

## Supplementary Materials and Methods

### MATERIALS AND METHODS: *IN VIVO*

#### Pharmacokinetic study in rodents

Pharmacokinetics of M102 was evaluated via single-dose subcutaneous and oral administrations of M102 in male C57Bl/6 mice. For subcutaneous administration, nine male C57BL/6J mice were administered with a single dose of 5 mg/kg M102. Blood samples were collected from each animal at 0.0833, 0.167, 0.25, 0.5, 1, and 2 hr post dose. For oral administration, a total of 18 male C57Bl/6 mice were dosed at 10 mg/kg, with three animals per blood sampling time point, which was determined as 0.25, 0.5, 1, 2, 3, and 6 hours post dose. Following blood collection from individual animals, plasma samples were extracted and processed, and then analysed by an LC-MS/MS bioanalytical method to determine the M102 plasma concentrations.

To investigate the brain exposure of M102, additional pharmacokinetic studies were also conducted in Spray-Dawley rats. Single dose oral administration of M102 at 12 mg/kg was given to two groups of male rats (N= 3/group). Plasma and brain samples were collected at 30 or 60 min post dose for two individual groups. Bioanalysis were conducted for plasma and brain samples to determine the M102 concentrations via LC-MS/MS.

Data acquisition was performed by LabSolutions version 5.89 Software (Shimadzu). All concentration data were reported with 3 significant figures. Data statistical analysis was performed using Microsoft Excel® software. The PK parameters of M102 were calculated using a non-compartmental approach with Pharsight® WinNonlin® v5.2 software package.

#### General toxicology study in rats and non-human primates (NHPs)

GLP general toxicology studies in rats and NHPs were conducted at WuXi AppTec (Suzhou, China) to evaluate potential toxicity of M102 when administered once daily for 28 days by oral gavage and to assess the reversibility, persistence, or delayed occurrence of toxic effects following a 14-day recovery period.

M102 at doses of 25, 50 or 75 mg/kg/day by oral gavage was investigated in the study of each species, based on previous non-GLP general toxicology study results. Criteria for evaluation included: viability (morbidity/mortality), clinical observations, body weight, food consumption, ophthalmology, clinical pathology (hematology, coagulation, serum chemistry, and urinalysis), toxicokinetics (TK), gross pathology, organ weights, and histopathology. Based on the study outcomes, no observed adverse effect levels (NOAELs) in both rats or NHPs were determined.

The protocol and additional amendments involving the care or use of animals involved in this study have been reviewed and approved by WuXi AppTec Institutional Animal Care and Use Committee (IACUC) prior to the initiation of study procedures. In addition, an onsite staff veterinarian monitored the study for animal welfare issues.

Toxicokinetic (TK) analysis of M102 plasma concentration-time data was performed at the Testing Facility. TK parameter values, including the maximum plasma concentrations (C_max_), the time to reach the maximum concentrations (T_max_), and the area under the plasma concentration vs. time curve (AUC) from time zero to 24 hours post-dose (AUC_0-24h_), were determined using a validated WinNonlin program. Values that were below the lower limit of quantification (BLQ) were set to zero in the calculation of the mean concentrations. Male and female TK data were analyzed separately. An audited TK report was generated and is attached to the Final Report as an appendix. Safety margin calculations were performed based on the resulting TK data and available preclinical *in vivo* PK data in mice, using the WinNonlin program.

#### Murine therapeutic study design

Transgenic female mice were block randomised into the different dose groups. Time points for behavioral tests were determined through analysis of previous characterisation of the models. The number of mice per group was determined through power analysis based on the standard deviation seen in the read-outs in previous experiments. Power analysis was based on our previously published model characterisations (*32, 46*).

#### Dosing

The oral formulation of M102 was in 10% HP-b-CD in water, which was filter sterilised before addition of M102. The subcutaneous formulation of M102 was in 1% ascorbic acid (Sigma-Aldrich), 0.05% sodium metabisulfite (Sigma-Aldrich) and 0.9% saline adjusted to pH 3.5. This was filter sterilised before addition of M102. M102 was weighed out into bijous and protected from light. On the morning of dosing M102 was dissolved in the appropriate vehicle for either oral or subcutaneous dosing to create the correct concentrations. Formulation of M102 was aided by vortexing and heating. Mice were dosed at 10 ml/kg and body weight was recorded daily for calculating doses.

#### Rotarod

Animals were tested once per week (TDP-43^Q331K^ mice) or twice per week (SOD1^G93A^ mice) on an accelerated rotarod (ACCELER rota-rod for mice, Jones & Roberts). The rotarod protocol used accelerated from 4 to 40 rpm over 5 minutes. Before testing started, mice were acclimatised to the equipment for three consecutive days with two runs per day. On test days, the mice were run twice on the rotarod protocol with a rest period in between and the time taken to fall off the equipment was recorded. Analysis was carried out using the best performance run on each testing day.

#### Catwalk gait analysis

Gait analysis was conducted using the Catwalk gait analysis system (Noldus), which consists of a glass plate and a green LED inside that is internally reflected when paws make contact with the surface and a camera mounted underneath. Mice ran backwards and forwards on the glass until 6 straight continuous runs were collected. Analysis was carried out with Catwalk software (Noldus version 7.1) with further analysis carried out using excel (Microsoft) and Graphpad prism.

#### Electrophysiology: compound muscle action potential (CMAP) and repetitive stimulation

Mice were anesthetised with 5% isoflurane and 4L/min oxygen in an induction box and then maintained with a nose cone under 2% isoflurane and 0.5L/min oxygen, which was adjusted as needed per mouse to keep breathing regular. The left hind limb of the mouse was then shaved and the fine fur was removed using hair removal cream (Veet). After loss of pedal reflex, a grounding electrode (Ambu Neuroline) was placed in the tail of the mouse, with the reference ring electrode being placed around the ankle covered in Ten20 nerve conductive paste (Weaver and Company) to ensure good contact with the skin. The recording ring electrode was then placed around the top part of the leg. The sciatic nerve was stimulated with a square input pulse with a 0.1ms duration, using a pair of twisted electrodes (Ambu Neuroline) that stimulated the hind limb. The CMAP amplitude was detected by the reporting electrode for the hind limb and the amplitude was recorded. The starting stimulating amplitude was approximately 5mV which was then increased gradually until the supramaximal response was determined for the CMAP, which was recorded for analysis. Repetitive stimulation was carried out by stimulating the muscles at the supramaximal amplitude for 10 stimuli at 10Hz to determine if the response declined over time. The amplitude of each response was recorded and analysed.

#### Tissue collection

At the end of the studies, mouse tissues were collected for histological analysis through perfusion or for protein and RNA analysis through a method of snap freezing and exposure to RNA later. Mice for perfusion were overdosed with pentobarbital and, after loss of the pedal reflex, were perfused with PBS followed by 4% paraformaldehyde. Brain and spinal cord were removed and stored in 4% PFA overnight before being stored in PBS.

#### Preparation of lumbar spinal cord sections and histology

A 12mm segment of lumbar enlargement was collected from the PFA fixed animals and embedded in two halves in paraffin wax. Sections were cut at a thickness of 10 μm and placed onto glass slides in a serial manner in blocks of 5 slides. Paraffin was removed from the sections, and they were rehydrated by washing in 100% xylene and a decreasing concentration gradient of ethanol.

#### Nissl staining

Slides were incubated with 0.1% cresyl violet for 15 minutes and then washed in water and incubated in acetic acid for 4 – 8 seconds followed by 100% ethanol for 1 minute and 100% xylene for 1 minute. These were then mounted and imaged using a nanozoomer digital slide scanner (Hamamatsu). Motor neuron counting was performed blinded and the criteria for counting of motor neurons were: cell body size greater than 20 μM in any axis excluding processes; location in the spinal cord ventral horn; and the presence of a nucleus and nucleolus.

#### IBA1 and GFAP staining

Slides underwent antigen retrieval at pH9 using an antigen access unit (a. Menarini Diagnostics) for 30 minutes at 125°C with 20 psi. Slides were blocked with 5% BSA and 0.25% triton for 20 minutes. Primary antibodies were added at a 1 in 500 dilution in a solution of 1% BSA and 0.25% triton PBS and incubated at 4°C overnight. Primary antibodies were anti-GFAP raised in chicken (Abcam, ab4674) and anti-Iba1 raised in rabbit (GeneTex, GTX1000042). Slides were washed with PBS and secondary antibodies were added at a 1 in 1000 dilution in a solution of 1% BSA in PBS. Secondary antibodies were 488 anti-chicken (Life Technologies, A11039) and 555 anti-rabbit (Thermo Fisher, A27039). Slides were incubated at room temperature in the dark for 1.5 hours. Slides were mounted with hardset Vectashield with DAPI (Vector Laboratories). Fluorescent images were captured using the INCELL 2000 system (GE). Images of the ventral horns of each spinal cord section were captured at 60 X objective magnification using DAPI (excitation 350 nm and emission 455 nm), FITC (excitation 490 nm and emission 525 nm) and Cy3 (excitation 543 nm and emission 605 nm) filters.

#### RNA extraction

RNA was extracted from brain and spinal cord using the RNeasy lipid mini kit (Qiagen, 74804) following the manufacturer’s protocol. RNA concentration and purity were checked using a nanodrop ND-1000 spectrophotometer (Thermo Scientific) by measuring absorbance at 230, 260 and 280 nm.

#### cDNA synthesis

cDNA was synthesised from RNA by firstly digesting DNA from 2000ng of RNA using RNase-free DNase and DNase buffer (Roch Diagnostics). Random hexamer primers and deoxyribonucleotide triphosphates were added to each reaction and incubated at 75°C for 5 minutes to denature RNA, and samples were placed immediately on ice. Reverse transcriptase was added (Invitrogen, 28025-013) to all samples and these were placed in a PCR machine with the following protocol: 25°C for 10 minutes, 42°C for 1 hour, 85°C for 5 minutes.

### RT-qPCR

RT-qPCR was carried out on cortex and spinal cord cDNA using optimised primers for targets that were downstream of NRF2 or HSF1. *Gapdh* was used as an endogenous control and data were analysed using the delta-delta method to gain relative gene expression to vehicle dosed samples.

### MATERIALS AND METHODS: *IN VITRO*

#### Immunohistochemistry to demonstrate a marker of oxidative stress in human postmortem tissue

Formalin-fixed, paraffin-embedded (FFPE) human CNS post-mortem tissue obtained from the Sheffield Brain Tissue Bank was used to investigate the level of nucleic acid oxidation in ALS patients versus controls using 8-OHdG as a marker of oxidative stress. Using a case control structure, the cohort consisted of ALS samples with age matched controls of: (i) frontal cortex (Fcx); (ii) motor cortex (Mcx) and (iii) spinal cord (SC). Immunohistochemistry was carried out on sections from 30 FFPE blocks derived from 10 individuals: 3 controls, 6 sALS, 2 C9-ALS and 2 C9 ALS-FTD (**Suppl. Table 2**).

FFPE blocks were sectioned at 6 μm and immunohistochemistry was performed using a standard ABC method (Vector Laboratories, Peterborough, UK) and the signal visualized using 3,3ʹ-diaminobenzidine. A summary of the primary antibody and the conditions, including antigen retrieval, is shown in **Suppl. Table 6**. Negative controls consisted of sections incubated with isotype controls or with omission of the primary antibody.

#### Detection of oxidative stress in human CSF by enzyme linked immunosorbent assay (ELISA)

The level of expelled fractions of oxidation in the CSF of ALS patients and controls was determined by ELISA to detect the oxidised RNA fraction. RNA oxidative stress has been reported to be directly associated with neurodegenerative pathology. The level of RNA oxidation was measured in 26 human CSF samples: 14 ALS patients and 12 controls (**Suppl. Table 3**) using the Cell Bio-labs Oxiselect RNA 8-OHdG ELISA kit (Catalog Number STA-325). The unknown 8-OHG CSF samples or 8-OHdG standards were first added in absolute concentration to an 8-OHdG/BSA conjugate preabsorbed EIA plate. Following a short incubation, an anti-8-OHdG monoclonal antibody was added, followed by an HRP conjugated secondary antibody. The quantity of 8-OHdG in the unknown sample was determined by comparing its absorbance with that of a known 8-OHdG standard curve. The kit has an 8-OHdG detection sensitivity range of 300 pg/mL to 40 ng/mL.

#### Immunofluorescence imaging of iAstrocytes

After fixation in 4% PFA for 10 min at RT, cells were washed (2x 5 min) in PBS (137 mM NaCl, 2.68 mM KCl, 10.14 mM Na2HPO4, 1.76 mM KH2PO4), blocked and permeabilised in 5% horse serum (DAKO) with 0.5% triton X-100 (AppliChem) for 30-60 min at RT, and incubated in primary antibody ON at 4°C (details of the primary antibodies in **Suppl. Table 7**). The cells were then washed (1x 5min in 0.2% Tween-20, Sigma, and 2 x 5min in PBS) and incubated in 1:1000 dilution of secondary antibody for 30-60min at RT (details of the secondary antibodies in **Suppl. Table 7**). Finally, the cells were stained with Hoechst 33342 (1:6000, Life Technologies) for 5-10 min at RT before washing (as previously) and then stored at 4°C. Imaging was performed with the OPERA Phoenix high-content imaging system (Perkin Elmer). Harmony 4.9 Analysis Software (Perkin Elmer) was used to analyse and quantify the intensity of immunostaining.

#### Immunoblotting of i-Astrocyte lysates

Protein extraction was performed by lysing the cells in IP lysis buffer (150 mM NaCl, 50 mM HEPES, 1 mM EDTA, 1 mM DTT, 0.5% (v/v) Triton X-100, PIC, pH 8.0) containing protease and phosphatase inhibitors, for 15 min on ice, followed by a centrifugation at 17,000g for 5 min at 4°C. Alternatively, the cells were lysed with RIPA buffer with 10% proteinase inhibitor and sonicated using the Soniprep 150 (MSE) for 10 seconds at 25% amp. before centrifuging as previously. The supernatant was removed and discarded, and this last step was repeated to wash the cell pellet, ensuring that all RIPA-soluble protein had been removed before the addition of urea. The supernatant was replaced with 10 μl of urea buffer and the sample was pipetted a few times to solubilise RIPA-insoluble material. The protein lysates were quantified using a bicinchoninic acid (BCA) assay (BCA assay kit, ThermoFisher Scientific), following manufacturer’s instructions, or by Bradford assay.

A total of 20-30 µg of protein mixed with 4x Laemmli buffer (228 mM Tris-HCl, 38% (v/v) glycerol, 277 mM SDS, 0.038% (v/v) bromophenol blue, 5% (w/v) β-mercaptoethanol pH 6.8) was denaturated at 95°C for 5 min and loaded into an SDS-polyacrylamide gel (12% resolving gel and 5% stacking gel). Resolving gel: 3.5 ml of dH2O2, 1.5 ml of 30% acrylamide, 2.5 ml of resolving buffer (1.5M Trizma®, 13.9mM SDS, pH 8.8, filtered), 50 µl of 10% APS, 20 µl of TEMED. Stacking gel: 5.8 ml of dH2O2, 1.7 ml of 30% acrylamide gel, 2.5 ml of stacking gel (0.5M Trizma®, 13.9 mM SDS, pH 6.8, filtered), 50 µl of 10% APS, 20 µl of TEMED. The SDS-polyacrylamide gel, assembled into a Mini-PROTEAN® Tetra Vertical Electrophoresis Cell (BioRad) filled with running buffer (25 mM Tris, 3.5 mM SDS, 20 mM glycine), was run for approximately 1.5h at 150-180V using a PowerPac Basic (BioRad). Then, the protein was transferred from the gel onto a nitrocellulose membrane (GE Healthcare) pre-soaked in transfer buffer (47.9 mM Tris, 38.6 mM glycine, 1.38 mM SDS, 20% (v/v) methanol), using the semi-dry Biometra FastblotTM transfer system (Analytik Jena), at 0.15A/gel for 1h.

The nitrocellulose membrane was blocked for 1h in 5% (w/v) milk in tris-buffered saline with Tween-20 (TBST) (20 mM Tris, 137 mM NaCl, 0.2% (v/v) Tween® 20, pH 7.6), at RT. After blocking, the membranes were incubated in primary antibody (**Suppl. Table 8**) diluted in blocking buffer, ON at 4°C. After 3x 5 min washes in TBST, the membrane was incubated in secondary antibody (anti-rabbit IgG HRP, or anti-mouse IgG HRP, Promega) diluted in blocking buffer for 1 h at RT. The membrane was then washed as previously, incubated in ECL for 1 min and scanned using the Odyssey® Fc Imaging System from LI-COR Biosciences.

#### Genome-wide RNA analysis of stalled protein synthesis (GRASPS) to detect translating mRNAs (71)

##### RNA extraction and ribosomal purification

RNA was extracted in Buffer A (250 mM sucrose, 5 mM KCl, 50 mM Tris-HC) containing proteinase inhibitor, 2 mM PMSF and 0.16 U/μl RNAse inhibitor. A final concentration of 0.7% v:v NP-40 from a 10% stock solution was gently mixed into the lysate. The lysate was incubated on ice for 5 minutes, mixed and incubated on ice for another 5-10 minutes. On a 6 well plate, the lysate was UV-irradiated on ice at 0.3 J/cm² and then centrifuged at 750 g for 10 minutes at 4°C to pellet the nuclear fraction. This nuclear pellet was discarded, and the supernatant was centrifuged at 12,500g for 10 minutes at 4°C to pellet the mitochondrial fraction. The supernatant, (post-mitochondrial (PMT) fraction) was transferred to a new cold Eppendorf tube and 4 M KCl was added to give a final concentration of 0.5M KCl. 1 ml of sucrose cushion was added to the bottom of clean, cold TLA100 centrifuge tubes. The PMT fraction was made up to 1 ml with Buffer B (250 mM sucrose, 500 mM KCl, 50 mM Tris-HCl, 5 mM MgCl2) and 900 μl of the 0.5 KCl-adjusted PMT fraction was slowly dispensed on top of the sucrose cushion and the tubes were balanced precisely within 0.01 g of each other before centrifuging at 250,000 g/75,000 rpm for 2 hours at 4°C in the TL-100 benchtop ultracentrifuge (Beckman). The ribosome pellet was quickly washed with cold DEPC water before resuspending in 250 μl of ribosome resuspension buffer (RRB). 0.16 U/μl Ribosafe RNAse inhibitor and 100 μg/ml proteinase K were added to the RRB and this was incubated for 30 minutes at 37°C with a pulse vortex every 5 minutes. After incubation, 10 mM EDTA and 50 mM NaAc was added and the RRB was vortexed. 750 μl of PureZOL RNA isolation reagent was added and left for 10 minutes at room temperature before the RNA was extracted using the Direct Zol RNA Miniprep Plus kit.

##### mRNA purification

The mRNA was isolated using the NEB Next® Poly(A)+ mRNA Magnetic Isolation Module. 15 μl of Oligo d(T)25 beads were dispensed into 0.2 ml tubes and the beads were washed twice with 100 μl of 2 x RNA binding buffer to ensure removal of the supernatant. An equal volume (50 μl) of 2 x RNA binding buffer and sample RNA were added to the beads. The samples were incubated at 65°C for 5 minutes and immediately placed on ice for 2 minutes. The samples were incubated at room temperature for 5 minutes before they were placed on a DynaMagTM-2 magnet (Life Technologies) for 2 minutes. The supernatant was removed and kept on ice. Previous work investigating this technology had incorporated an additional binding step which increased the yield of ribosomal poly(A)+ RNA. The beads were washed twice with 200 μl of wash buffer and placed on the magnetic rack for 2 minutes. The beads were then stored on ice while the initial saved supernatant was heated to 65°C for 5 minutes and directly on ice for 2 minutes. The supernatant was then re-added to the beads and the binding of the poly(A)+ RNA was repeated as above. After the supernatant was removed and kept on ice, the beads were washed twice with 200 μl wash buffer. After ensuring the total removal of the wash buffer, 50 μl of Tris-buffer was added to the beads. The samples were incubated at 80°C for 2 minutes then immediately left at room temperature. 50 μl of RNA binding buffer was then added to the beads and the samples were incubated at room temperature and inverted every few minutes before placing the tubes on the magnetic rack and removing the supernatant. The beads were washed twice again in 200 μl of wash buffer. The poly(A)+ mRNA was eluted from the beads by adding 20 μl of Tris-buffer, incubating samples at 80°C for 2 minutes and then placing on the magnetic rack to transfer the supernatant to a new nuclease-free PCR tube. The sample was then added to the beads and the binding of the poly(A)+ was repeated as described above. After this incubation, the tubes were placed on the magnetic rack to remove the supernatant. The mRNA was again eluted from the beads through the addition of 20 μl of Tris-buffer and the process described above. The quantity of purified mRNA was assessed using the NanoDrop Spectrophotometer ND-1000 (Labtech International).

##### Ribosomal RNA depletion

3 μl of the RNA/probe Master Mix (1 μl of NEBNext rRNA depletion solution, 2 μl probe hybridisation buffer) was added to 12 μl of RNA sample and pipetted up and down 10 times. The samples were briefly spun in a tabletop centrifuge and immediately placed in the G-STORM thermocycler (LABCARE) with a heated lid of 80°C running the following program: 95°C for 2 min, 22°C for 5 min. The samples were spun down briefly and placed on ice while the RNase H Digestion Master Mix (2 μl NEBNext RNase H, 2 μl of RNase H reaction buffer, 1 μl nuclease-free water). 5 μl of the above mix was added to each sample and was mixed thoroughly with a pipette. The samples were briefly spun and immediately placed in a thermocycler with a heated lid of 40°C and incubated at 37°C for 30 minutes. After incubation, the samples were spun down briefly again and placed on ice while the DNase I Digestion Master Mix (5 μl DNAse I reaction buffer, 2.5 μl DNase I, 22.5 μl nuclease-free water). 30 μl of the above mix was added to each sample and thoroughly mixed with a pipette. The samples were briefly spun and immediately incubated at 37°C for 30 minutes in the thermocycler. After incubation, the samples were spun down again and placed on ice. A 1 ml aliquot of NEBNext® RNA Sample Purification Beads was vortexed into suspension and 110 μl of this suspension was added to each sample. The samples were then thoroughly mixed with a pipette and incubated on ice for 15 minutes. The tubes were placed on a magnetic rack to separate the beads from the supernatant. After 5 minutes, the supernatant was removed and discarded, and the beads were washed twice with 200 μl of freshly prepared 80% ethanol; beads were incubated in ethanol for 30 seconds between washes. The tubes were briefly spun to remove any excess ethanol. The beads were then air dried for 5 minutes while the tubes were left on the magnetic rack with the lids open. The samples were eluted while the beads were still dark brown and glossy looking; if the beads were over-dried this would impact the yield. 8 μl of nuclease free water was added to the beads, thoroughly mixed, and incubated on ice for 2 minutes to elute the RNA sample from the beads. The tubes were placed on the magnetic rack for 5 minutes to sediment the beads and the RNA sample was transferred to a fresh 0.2 μl Eppendorf.

#### RNA Sequencing

RNA sequencing was performed at the Centre of Genomic Research, University of Liverpool, using a dual-indexed, strand-specific RNA-Seq library from the submitted poly(A)+ enriched RNA sample using the NEB Next Ultra Directional RNA library preparation kit. Samples were sequenced on lanes of the Illumina Nova Seq using S4 chemistry (paired-end, 2 x 150 base pair sequencing) which generated approximately 2500 M clusters per lane. RNA sequencing data were quality controlled at the centre by removal of any low-quality bases and sequencing adapters.

##### Quality control

Bcbio is a python toolkit providing best-practice pipelines for fully automated high throughput sequencing analysis (ttps://bcbio-nextgen.readthedocs.io/en/latest/). The bcbio pipeline was used to merge all 5-6 FastQ files per sample together for quality control and further processing. Inspecting the GC-content histograms from FastQC https://www.bioinformatics.babraham.ac.uk/projects/fastqc/) revealed a bimodal distribution for some samples attributed to ribosomal contamination. Therefore, the bbsplit tool was used to remove any residual rRNA sequences prior to alignment.

##### Sequence alignment and differential expression

*Salmon* (https://salmon.readthedocs.io/en/latest/) was used to quantify the expression of transcripts using the RNA sequencing data and to perform an inference step to estimate the relative abundance of all the known transcripts without aligning the reads. The genecode set of transcripts (v28) was used for quantification, giving a matrix of transcript-level quantifications for each sample. The tximport Bioconductor package was used to aggregate the counts to the gene-level and these counts were imported into the DESeq2 Bioconductor package for quality control and analysis. Visualisation of the raw count distributions and unsupervised analysis prompted the removal of two outlier subjects from further analysis (control 155 and C9orf72 patient 183). The DESeq2 differential expression method incorporates gene-length and library size correction as part of the model. Prior to visualisation in other analysis tools, the RNA sequencing data were normalised by the FPKM (fragments per kilobase per million reads) method. Lists of genes that were statistically differentially expressed were identified by applying a p-value <0.05 and absolute log2 fold-change >1.5. The gene lists were inputted into DAVID Functional Annotation Bioinformatics Microarray Analysis programme (https://david.ncifcrf.gov/). The list of Gene Ontology Term “Biological Process” (GO-BP) terms were exported from DAVID to categorise differentially expressed genes (DEG). To visualise group segregation based on DEGs, principal component analysis (PCA) plots and heatmaps were generated using Qlucore Omics Explorer software (Qlucore).

## Supplementary Figures and Tables

**Supplementary Figure 1:**
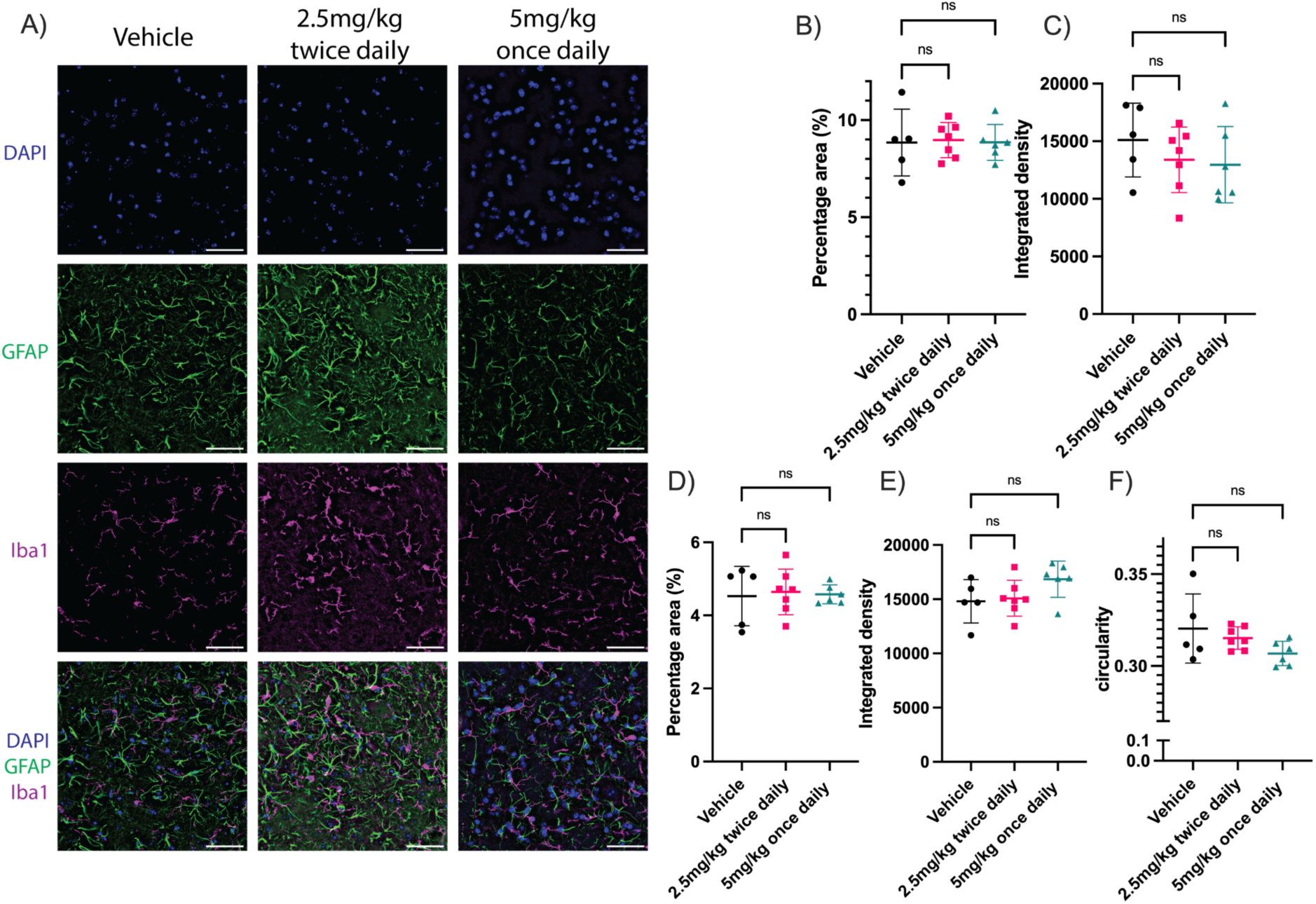
M102 shows no alteration of astrocyte or microglia staining in the spinal cord of TDP-43^Q331K^ mice. A) Representative images of lumbar spinal cord from 6 month old mice showing individual and composite images for staining with DAPI, GFAP and Iba1. Scale bar =50μm. B) percentage area staining and C) integrated density of GFAP staining. D) Percentage area staining, E) integrated density and F) circularity of Iba1 staining. One-way ANOVA with Dunnett’s multiple comparisons. Data shown as individual points with mean +/- SD (N=5-7 mice per group).

**Supplementary Figure 2:**
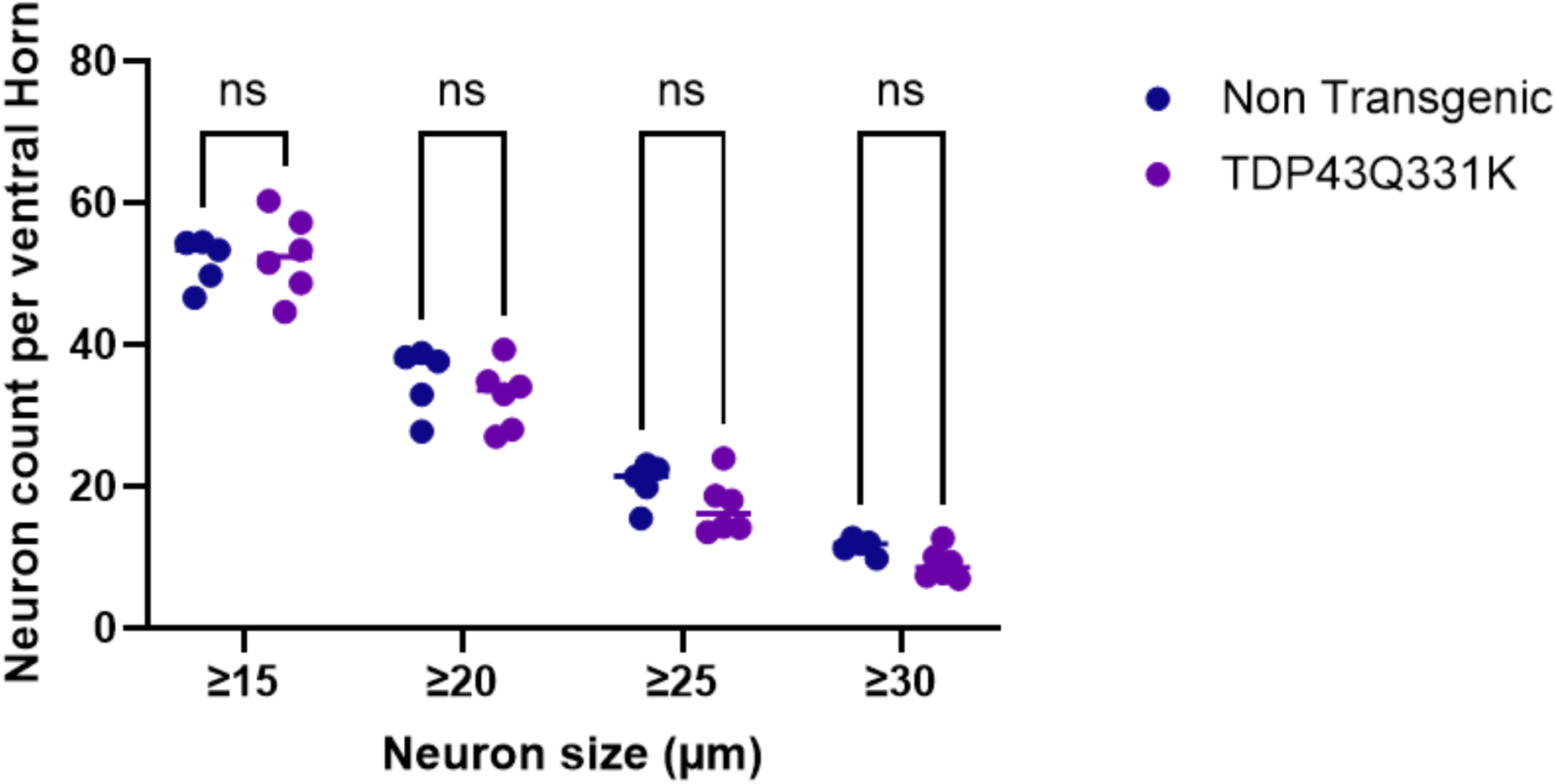
Lumbar ventral horn motor neuron counts in TDP43Q331K transgenic and non-transgenic littermates. Motor neurons were counted as described in the supplementary methods in size bins ranging from >15 to >30 microns (single chord length within cytoplasm) from spinal cord tissue collected from 5 month old mice. No significant differences were seen between the TDP43Q331K transgenic mice and non-transgenic littermates. Two-way ANOVA overall p=0.36 for effect of genotype

**Supplementary Figure 3:**
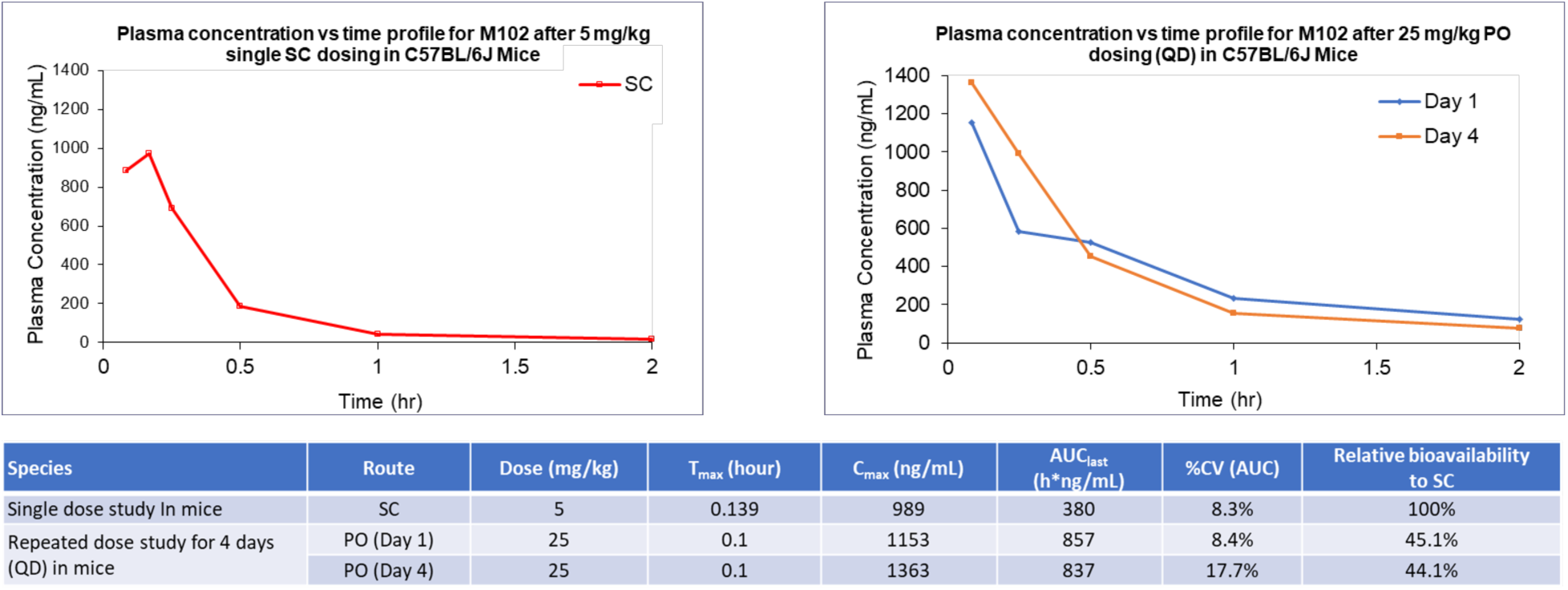
Oral gavage dose of 25mg/kg is bioequivalent to 5mg /kg subcutaneous dose in mice. A) Pharmacokinetic study of M102 following single subcutaneous (sc) dose at 5mg /kg or B) oral dosing for 1-4 days at 25 mg/kg/day PO (QD). Data show plasma concentrations as determined by HPLC-MS, which was equivalent to 5mg/kg/day SC, assuming 20% relative bioavailability In the multiple dose study. Day 4 showed consistent PK profile to Day 1 Pharmacokinetic profile of M102 dosed at 5mg/kg SC and 10 mg/kg PO. Multiple Oral Dose Study Showed Consistent Exposure.

**Supplementary Figure 4:**
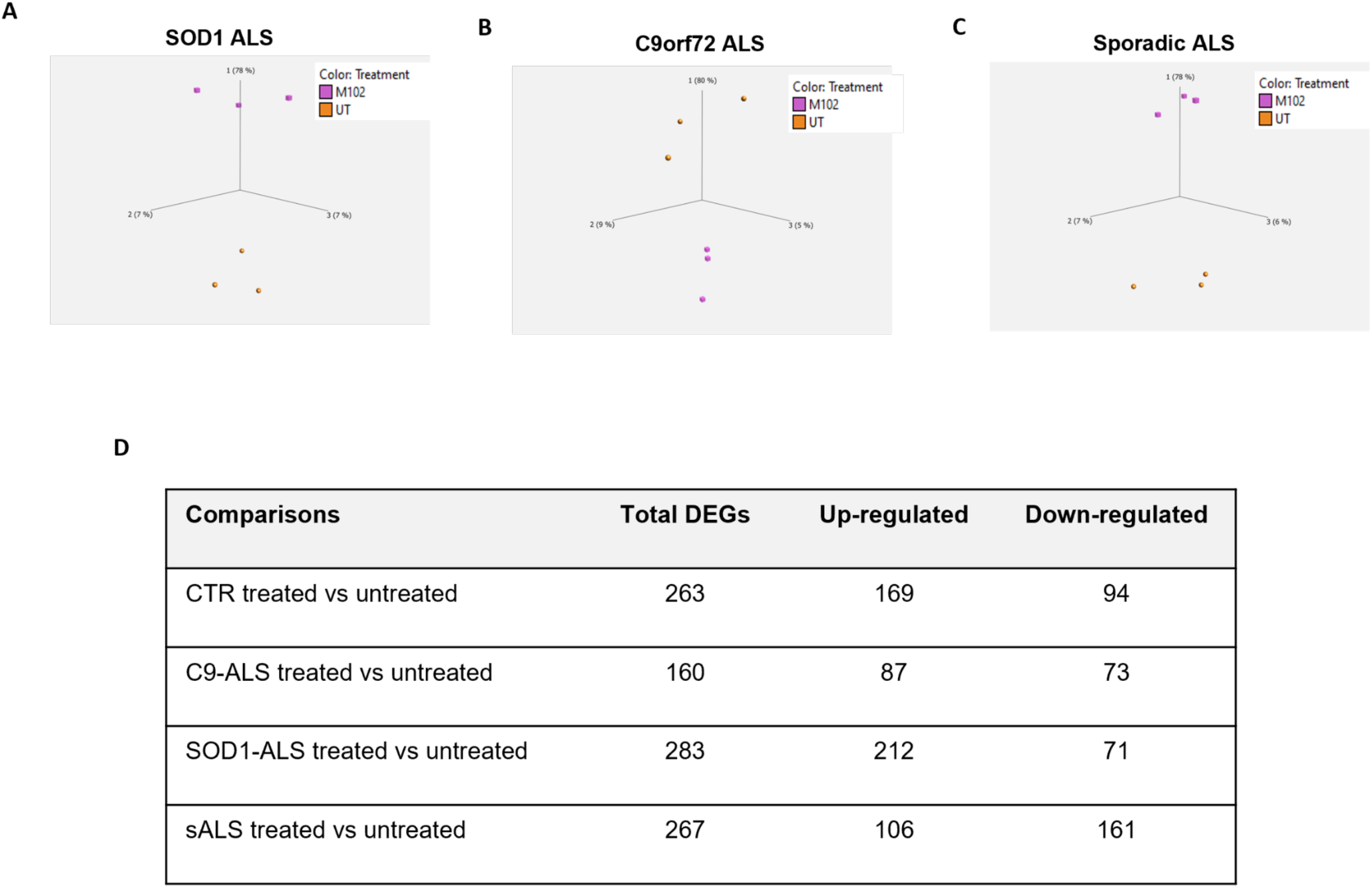
RNA sequencing from iAstrocytes treated with 10 μM M102 or DMSO for 48h reveal genotype specific transcriptomic changes in response to M102 treatment. These transcriptomic changes are observed across A) SOD1, B) C9orf72, and C) sporadic ALS cases, suggesting that M102 has the potential to be beneficial for both familial and sporadic ALS patients. D) Summary of the DEGs identified between treated and untreated iAstrocytes from healthy controls, C9-ALS, SOD1-ALS and sALS cases.

**Supplementary Table 1:**
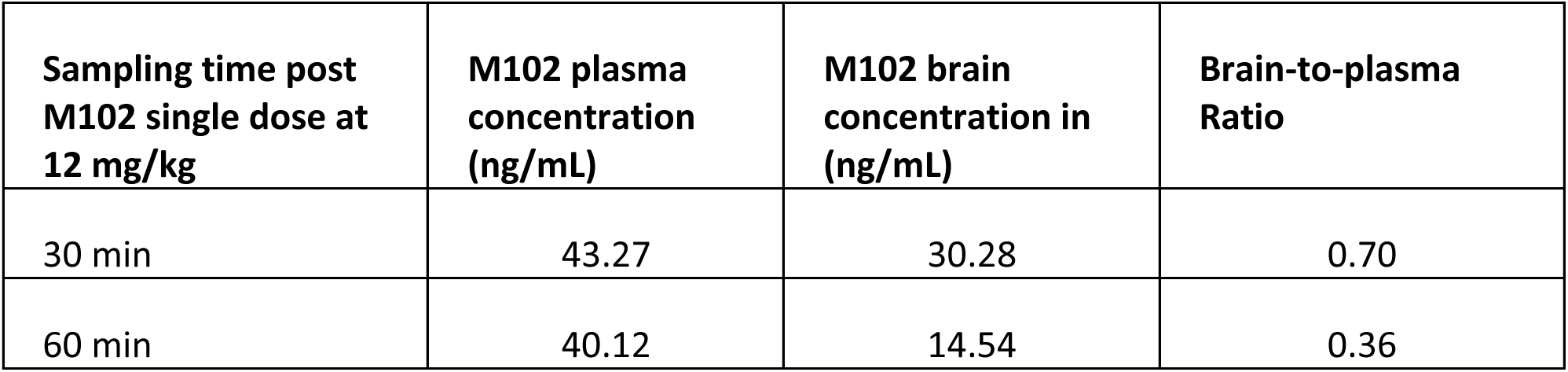
Brain penetration of M102 in male Sprague-Dawley rats.

| **Sampling time post**<br>**M102 single dose at**<br>**12 mg/kg** | **M102 plasma**<br>**concentration**<br>**(ng/mL)** | **M102 brain**<br>**concentration in**<br>**(ng/mL)** | **Brain-to-plasma**<br>**Ratio** |
| --- | --- | --- | --- |
| 30 min | 43.27 | 30.28 | 0.70 |
| 60 min | 40.12 | 14.54 | 0.36 |

**Supplementary Table 2:** Post-mortem CNS tissue used for 8-OHG staining.

| **Case ID** | **Gender** | **Age** | **PMD** | **Diagnosis** | **Comments** |
| --- | --- | --- | --- | --- | --- |
| 98-07 | M | 67 | 63hrs | Hepatocellular carcinoma |  |
| 10-96 | F | 63 | 31hrs | Carcinoid Tumor |  |
| 07-70 | M | 26 | Not available | Epilepsy (sudden death) | No structural pathologic abnormality |
| 05-64 | M | 66 | 96hrs | sALS |  |
| 08-24 | F | 63 | 36hrs | sALS |  |
| 08-96 | F | 69 | 40 hrs | sALS |  |
| 14-11 | M | 51 | 40hrs | sALS |  |
| 05-42 | M | 66 | 9hrs | sALS |  |
| 59-09 | F | 80 | N/A | sALS |  |
| 69-09 | M | 48 | 23hrs | fALS C9orf72 |  |
| 83-10 | M | 79 | 13hrs | fALS C9orf72 |  |
| 41-04 | M | 79 | 13hrs | ALS+FTD C9orf72 |  |
| 66-08 | F | 59 | 28hrs | fALS+FTD C9orf72 |  |

**Supplementary Table 3:** CSF samples used for the 8-OHG ELISA.

| **Sample** | **Disease** | **Gender** | **Date of birth** | **Date of collection** | **Age at collection** |
| --- | --- | --- | --- | --- | --- |
| 3624 | Sporadic ALS | Male | 07/02/1962 | 20/11/2015 | 53 |
| 3629 | Sporadic ALS | Male | 29/04/1946 | 10/12/2015 | 69 |
| 427 | Sporadic ALS | Male | 05/09/1950 | 13/02/2014 | 63 |
| 3392 | Sporadic ALS | Female | 11/02/1951 | 20/11/2014 | 63 |
| 240 | Sporadic ALS | Male | 25/01/1952 | 21/03/2006 | 54 |
| 248 | Sporadic ALS | Male | 29/04/1940 | 27/03/2006 | 65 |
| 254 | Familial ALS<br>(mutation unknown) | Male | 03/10/1961 | 11/04/2006 | 44 |
| 303 | Sporadic ALS, MS | Female | 28/01/1943 | 10/10/2006 | 63 |
| 3575 | Sporadic ALS | Female | 26/11/1967 | 27/08/2015 | 47 |
| 3543 | Sporadic ALS | Male | 08/04/1953 | 06/07/2015 | 62 |
| 2307 | Familial ALS (M337V<br>TARDBP mutation) | Female | 15/06/1973 | 09/07/2015 | 42 |
| 3624 | Sporadic ALS | Male | 07/02/1962 | 20/11/2015 | 53 |
| 3629 | Sporadic ALS | Male | 29/04/1946 | 10/12/2015 | 69 |
| 3653 | Sporadic ALS | Male | 14/05/1975 | 17/02/2016 | 40 |
| C3631 | Healthy control | Male | 30/10/1959 | 15/12/2015 | 56 |
| C3634 | Healthy control | Female | 21/06/1950 | 01/02/2016 | 65 |
| C209 | Control:Headache | Female | 03/08/1955 | 06/06/2005 | 49 |
| C347 | Control:Depression | Male | 21/09/1970 | 09/02/2008 | 37 |
| C245 | Control:Facial weakness | Female | 03/07/1950 | 27/11/2014 | 64 |
| C3619 | Control:Headache | Female | 28/08/1987 | 17/11/2015 | 28 |
| C3606 | Control:Migraine | Female | 07/02/1970 | 14/10/2015 | 45 |
| C3549 | Control:Migraine | Female | 11/05/1978 | 22/07/2015 | 37 |
| C3548 | Healthy control | Female | 08/06/1989 | 15/07/2015 | 26 |
| C3576 | Healthy control | Male | 28/10/1973 | 01/09/2015 | 41 |
| C387 | Control: Atypical headache | Male | 28/03/1975 | 21/10/2008 | 33 |
| C355 | Control: Functional gait disorder | Male | 16/10/1959 | 09/09/2008 | 49 |

**Supplementary Table 4:** Cell lines used for in vitro experiments.

| **Cell line** | **Genetic remarks** | **Sex** | **Age at biopsy collection (years)** | **Ethnicity** | **Onset to death (months)** |
| --- | --- | --- | --- | --- | --- |
| CTR_1 | Non-ALS control | Male | 40 | Caucasian | - |
| CTR_2 | Non-ALS Control | Male | 31 | Caucasian | - |
| CTR_3 | Non-ALS control | Female | 64 | Caucasian | - |
| CTR_4 | Non-ALS control | Male | 68 | Caucasian | - |
| CTR_5 | Non-ALS control | Female | 69 | Caucasian | - |
| sALS_1 | sALS | Female | 61 | Caucasian | 21 |
| sALS_2 | sALS | Male | 29 | Caucasian | 90 |
| sALS_3 | sALS | Male | 47 | Caucasian | 72 |
| sALS_4 | sALS | Male | 59 | Caucasian |  |
| sALS_5 | sALS | Female | 85 | Caucasian | 25 |
| sALS_6 | sALS | Female | 80 | Caucasian | 29 |
| sALS_7 | sALS | Male | 54 | Caucasian | 37 |
| sALS_8 | sALS | Female | 58 | Caucasian | - |
| sALS_9 | sALS | Male | 56 | Caucasian | 49 |
| sALS_10 | sALS | Male | 65 | Caucasian | - |
| C9_1 | C9orf72 ALS | Male | 66 | Caucasian | 31.7 |
| C9_1 | C9orf72 ALS | Male | 50 | Caucasian | 27 |
| C9_3 | C9orf72 ALS | Female | 66 | Caucasian | 19.4 |
| SOD1_1 | SOD1 (A4V) | Female | 63 | Caucasian | - |
| SOD1_2 | SOD1 (A4V) | Female | 40 | Caucasian | - |
| SOD1_3 | SOD1 (D90A) | Male | 56 | Caucasian | - |
| SOD1_4 | SOD1 (D76Y) | Female | 42 | Caucasian | 81 |

**Supplementary Table 5:** List of GO pathways shared between ALS patient subgroups treated with M102.

| Gene Ontology Term | CTR p-value | C9ORF72 p-value | SOD1 p-value | sALS p-value |
| --- | --- | --- | --- | --- |
| Cellular response to organic cyclic compound | 4.10E-03 | - | 2.20E-04 | - |
| MAPK cascade | 2.70E-02 | - | 3.60E-02 | - |
| Morphogenesis of an epithelial fold | 4.40E-02 | - | - | 4.00E-02 |
| Cell adhesion | - | 3.40E-05 | 2.80E-02 | 4.30E-02 |
| VEGF pathway | - | 3.20E-02 | 1.80E-02 | 2.50E-02 |
| Inflammatory response | - | 6.20E-04 | 5.40E-05 | 3.80E-02 |
| Autophagic response | - | 2.40E-04 | 1.90E-03 | 2.70E-02 |
| Oxidation-reduction process | - | 5.60E-03 | 5.00E-02 | 3.70E-02 |
| Xenobiotic metabolic process | 5.60E-02 | - | 7.90E-04 | 4.40E-02 |
| Synaptic transmission, cholinergic | - | - | 3.70E-04 | 6.10E-03 |
| Collagen fibril organisation | - | - | 5.60E-03 | 6.20E-04 |

**Supplementary Table 6:** Primary antibody for immunohistochemistry of FFPE.

| **Antibody** | **Species** | **Dilution factor**<br>**and conditions** | **Antigen**<br>**retrieval** | **Supplier** |
| --- | --- | --- | --- | --- |
| 8-OHdG | Mouse<br>monoclonal | 1/400<br>1h RT | Pressure cooker<br>access<br>revelation (×10,<br>pH 6.5) | Abcam,<br>Cambridge UK |

**Supplementary Table 7:** Primary and secondary antibodies for immunocytochemistry of iAstrocytes.

| **Primary** | **Species** | **Dilution factor** | **Supplier** |
| --- | --- | --- | --- |
| 8-OHdG | Mouse monoclonal | 1:100 | Abcam, Cambridge UK |
| NRF2 | Rabbit polyclonal | 1:500 | Abcam |
| HSF1 | Rabbit polyclonal | 1:500 | Cell Signaling Technologies |
| mSOD1 | Mouse monoclonal | 1:100 | Medimabs |
| **Secondary** | **Species** | **Concentration** | **Supplier** |
| anti-chicken Alexa 488 | Goat | 2mg/ml | Invitrogen; A11039 |
| anti-rabbit Alexa 488 | Goat | 2mg/ml | Invitrogen; A11008 |
| anti-rabbit Alexa 568 | Donkey | 2mg/ml | Invitrogen; A10042 |
| anti-mouse Alexa 568 | Donkey | 2mg/ml | Invitrogen; A10037 |

**Supplementary Table 8:** Primary antibodies for immunoblotting of iAstrocytes.

| **Antibody** | **Species** | **Dilution and conditions** | **Supplier** |
| --- | --- | --- | --- |
| TDP43 | Rabbit polyclonal | 1:1000 | Proteintech |
| NQO1 | Rabbit polyclonal | 1:1000 | Abcam |
| HSF1 | Rabbit polyclonal | 1:1000 | Cell Signaling Technologies |
| B-actin | Mouse monoclonal | 1:10,000 | Abcam |

